# Geometric Network Analysis Provides Prognostic Information in Patients with High Grade Serous Carcinoma of the Ovary Treated with Immune Checkpoint Inhibitors

**DOI:** 10.1101/2021.06.10.447889

**Authors:** Rena Elkin, Jung Hun Oh, Ying L. Liu, Pier Selenica, Britta Weigelt, Jorge S. Reis-Filho, Dmitriy Zamarin, Joseph O. Deasy, Larry Norton, Arnold J. Levine, Allen R. Tannenbaum

## Abstract

**Purpose:** Network analysis methods can potentially quantify cancer disturbances in gene networks without introducing fitted parameters or variable selection. A new network curvature-based method is introduced to provide an integrated measure of variability within cancer gene networks. The method is applied to high grade serous ovarian cancers (HGSOCs) to predict response to immune checkpoint inhibitors (ICIs), and to rank key genes associated with prognosis.

**Methods:** Copy number alterations (CNAs) from targeted and whole exome sequencing data were extracted for HGSOC patients (*n* = 45) treated with ICIs. CNAs at a gene level were represented on a protein-protein interaction network to define patient-specific networks with a fixed topology. A version of Ollivier-Ricci curvature was used to identify genes that play a potentially key role in response to immunotherapy and further to stratify patients at high risk of mortality. Overall survival (OS) was defined as the time from the start of ICI treatment to either death or last follow-up. Kaplan-Meier analysis with log-rank test was performed to assess OS between the high and low curvature classified groups.

**Results:** The network curvature analysis stratified patients at high risk of mortality with p=0.00047 in Kaplan-Meier analysis. Genes with high curvature were in accordance with CNAs relevant to ovarian cancer.

**Conclusion:** Network curvature using CNAs has the potential to be a novel predictor for OS in HGSOC patients treated with immunotherapy.

## 1 Introduction

Facilitated by advances in genomic sequencing techniques and the ongoing development of highly curated protein-protein interactome (PPI) databases (e.g., Human Reference Protein Database (HPRD, [1, 2]), The Human Reference Interactome (HuRI, [3]), Search Tool for the Retrieval of Interacting Genes/ Proteins (STRING, [4])), we adopt a network approach to investigate biological features pertaining to overall survival (OS) in ovarian cancer (OC) based on copy number alterations (CNAs) in tumor tissues. The past decade has seen a large rise in the development of methods for analyzing large, complex networks, as exhibited by the rapidly growing literature. We draw on geometric notions to inform about the network structure, defined by evidence-based interactions provided by the PPI. Our network analysis methodology is unsupervised without fitting parameters or feature selection and is not constrained to the underlying topology alone. Indeed, since cancer has been demonstrated to exhibit functional robustness in connection to geometric properties of its network representation [5], we utilize Ollivier’s discrete notion of Ricci curvature on weighted graphs, referred to as *Ollivier-Ricci (OR) curvature* [6].

This focus of this paper is to introduce a geometric network method for cancer with the key application to high grade serous ovarian cancer (HGSOC). Biomarkers of response to immune checkpoint blockade in HGSOC remain largely unknown. Unlike non-small cell lung cancers and melanomas that exhibit increased immunogenicity due to high tumor mutational burden (TMB) [7, 8, 9, 10, 11], HGSOCs exhibit low TMB [12]. In virtually all cases, HGSOCs are a disorder of loss of function gene mutations (TP53) leading to CNAs, and subsequently resulting in over-expressed copy number in multiple genes including oncogenes (e.g., K-RAS, c-MYC, cyclin E and AKT protein kinase) commonly due to aneuploidy [13, 14]. The impact of these alterations on response to immunotherapy is unknown; furthermore, it is unlikely that individual pathway alterations would be strongly predictive. This manuscript develops a mathematical method that constructs a network of these gene pathways where each node (gene) is quantitated by CNAs and for each tumor, the changes in the architecture or connectivity of the network are measured by a parameter termed ***curvature*** of the edges of the network. Curvature measures the connectivity in the sense of feedback loops, and the copy number measures the abundance of each node and its projected impact upon the changes in the network architecture. (More rigorous details about this will be given in the Methods Section.) Nodal curvature may exhibit more variation than the CNAs, reflecting the integration of the gene copy numbers and the local impact of their alteration on the network. Thus, curvature has the potential to di↵erentiate responders from non-responders in patients treated with immune checkpoint inhibitors (ICIs) that could not be predicted from a single gene alone.

Curvature is a local measure of how a geometric object (e.g., curve, surface, space) deviates from being *flat* in the Euclidean sense. While the physical interpretation of curvature in 3-dimensional Euclidean space is a familiar concept, intuition for curvature as a rigorous mathematical concept is often elusive, as the mathematical theory is not bound by the same physical constraints. This allows for curvature to be generalized to continuous spaces of higher dimensions (classically, Riemannian manifolds), and even to discrete spaces (Figure 1).

**Figure 1:**
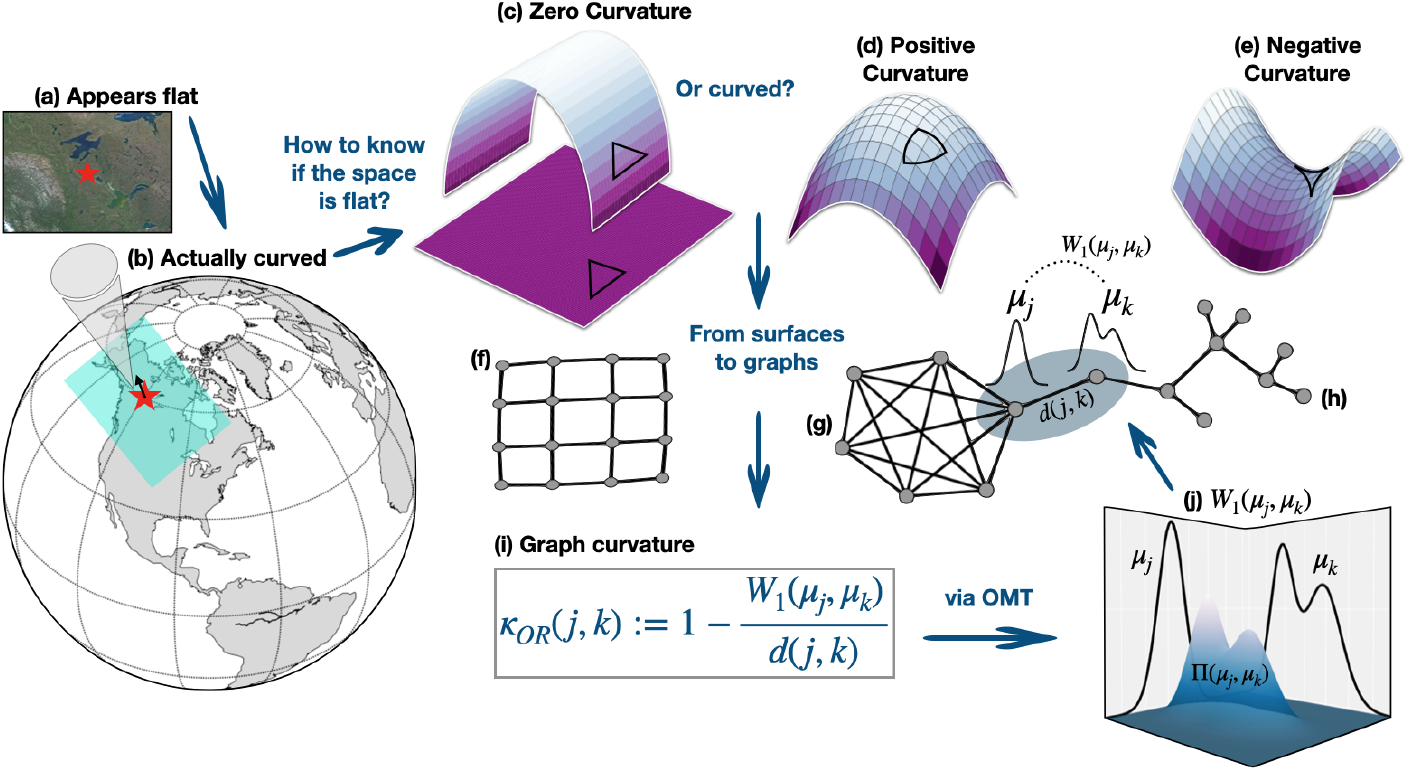
Curvature intuition on graphs. Curvature is an intrinsic property of a surface, and therefore does not depend on how it is situated in space. For example, (b) we know the Earth is curved, (a) even though it appears flat when standing on its surface. Similarly, (c) a plane that is bent into an arc still has zero curvature. The apparent curvature is merely due to how it is embedded in space. Examples of canonical surfaces with zero, positive and negative curvature are shown, respectively, in (c), (d) and (e). Geodesic triangles can be used to determine the curvature of a surface without specifying its embedding. Compared to (Euclidean) flat space (c), fat (d) and skinny (e) triangles are characteristic of positive and negative curved spaces, respectively. Going from smooth surfaces to graphs, (f) a grid is analogous to a surface with zero curvature while (g) many triangles (indicative of redundancies or feedback mechanisms) are characteristic of graphs with positive curvature and (h) tree-like topologies (indicative of diverging paths) are characteristic of graphs with negative curvature. (i) On a graph, curvature between two nodes *j* and *k* is characterized by the ratio of the transport distance *W*_1_(*μ*_*j*_, *μ*_*k*_) between distributions *μ*_*j*_ and *μ*_*k*_ (defined respectively on nodes *j* and *k*) and the underlying ground distance *d*(*j, k*) between the two nodes. The transport distance (*W*_1_) comes from the theory of optimal mass transport (OMT) and provides a *functional* distance between the nodes that accounts for the shape of the distribution and amount of shared neighbors. Curvature is positive (resp., negative) when the transport distance (i.e., information) between nodes is smaller (resp., larger) than the ground distance between them, reflecting the ease with which information is shared between nodes.

The mathematical construct, however, is not solely of abstract, theoretical value. The archetypical example is the curvature of space-time which was integral to Einstein’s theory of general relativity. Although perhaps less intuitive, the geometric insight that curvature provides is applicable to other physical phenomena. In particular, change in OR curvature [6] has a strong mathematical connection to changes in robustness via change in entropy. Note that we are using *change in curvature* in the sense as a difference in curvature Δ*κ* between networks. This is a remarkable result facilitated by the theory of optimal mass transport (OMT) attributed to Sturm, Lott, and Villani [15, 16]. The change in OR curvature has previously been used as an effective quantitative proxy for the qualitative notion of changes in robustness in various types of networks [5, 17]. In the present work, we employ curvature to predict patient survival and investigate primary components of functional robustness as well as to identify key genes contributing to functional dysregulation in HGSOC.

Various biomarkers including PD-L1 and the spatial distribution and composition of the immune microenvironment are being investigated in the context of response to ICI [12], but the present work focuses on extracting information from gene level information. It is becoming more apparent that the use of genomic data (e.g., mutations, gene expression, CNAs) with the corresponding functional network representation can provide more insights into understanding the underlying biology of cancer. Thus, graph-based tools may be more powerful for investigating complex genomic networks than methods that aim to analyze and quantify the data independently.

Genomic networks have a topology (i.e., a connectivity structure), but they also have a geometry, i.e., curvature, which gives a measure of their functional robustness. Graph curvature is intimately related to the number of invariant triangles, i.e., *feedback loops* at a given vertex, and the curvature between two vertices describes the degree of overlap between their respective neighborhoods [18]. Informally, graphs with positive curvature characteristically contain many triangles (redundant feedback loops), contributing to its functional robustness with respect to a damaged or deleted edge. The more neighbors two given nodes have in common (i.e., triangles), the easier it is for information to flow between them. By weighing the ease with which information can be transferred from one node to another against the ground distance between them, curvature provides a local measure of functional connectivity compared to ordinary measures of connectivity which identify hubs based on degree. We show not only that the total curvature of a network can be used to predict overall patient survival in OC, but it is also more effective than standard clinical parameters such as TMB.

Typically, the curvature is computed on a network using the standard hop distance (where every edge in a path connecting two nodes is treated as a hop) with node weights that are continuous in nature (e.g., gene expression). Here, we use a ***weighted hop distance*** derived from the data as the underlying graph metric, so the distance between two nodes depends not only on the topology, but on the likelihood of interaction as well. Using node weights assigned by (discrete) CNAs, we show that curvature may also be informative in the discrete data setting. Furthermore, we show that the network topology without any additional information may be used as a reference to identify potential key players responsible for the functional robustness, even when limited data is available, as demonstrated in this study. Top identified genes such as TP53, whose known aberrant functional behavior has been attributed as a leading influence in the development/progression of ovarian cancer [19], serve as validation for the proposed methodology.

Specifically, we create a shared topology, but with sample-specific gene interaction networks. The interactions are taken from the HPRD, where the protein interactions are assumed to serve as a proxy for the underlying gene interactions. We then supplement topology (i.e., connectivity) with sample-specific node weights taken to be the given copy number data. For each network, curvature is then computed at three scales: on edges, nodes, and the entire network. Analogous to Ricci curvature defined on tangent directions at a point on a Riemannian manifold and its contraction *scalar curvature* defined on the points of the manifold, the formulation of OR curvature is computed on all edges in the network and scalar curvature is computed on all nodes by contracting the OR (edge) curvature with the invariant distribution associated with the weighted network [6]. The *total curvature* of the network is then computed by contracting the scalar curvature to a single scalar. (See Eq. (9) for the precise definition.)

## 2 Methods

We start with a brief, informal discussion on curvature to build some intuition before introducing the formal description of curvature as it was used in this work (Figure 1).

The remarkable property of Gaussian curvature is that it is intrinsic to the surface and therefore independent of how the surface is embedded in 3-dimensional space. For example, the Earth appears flat when looking into the horizon, yet we know that the Earth is round. Determining a surface’s curvature by visual inspection alone can be very misleading, as the curvature may appear to change depending on one’s perspective. More generally, suppose we take our surface to be a sheet of paper lying flat on a desk. One would correctly guess that it has zero curvature. If the scenario is changed and the paper is bent into an arc, it may appear to have non-zero curvature. However, this apparent curvature is merely an effect of its *embedding* in space and is not intrinsic to the surface itself. Thus, a plane and a cylindrical arc are all examples of surfaces with zero Gaussian curvature while a sphere and a hyperbolic disc are examples of surfaces with positive and negative Gaussian curvature, respectively [20].

Rather than look at the surface as it is embedded in 3-dimensional space from the perspective of an outsider, the key is to treat the surface as the space itself. In that case, we can determine if the space is curved through the use of ***geodesics***, the curves of (locally) shortest length between two points. (Geodesics generalize the straight line in Euclidean space.) One way to tell if the space is curved is to sum up the interior angles of a geodesic triangle. Geodesic triangles on a surface with positive (resp., negative) Gaussian curvature are *fat* (resp., *skinny*) compared to the triangle in Euclidean space. Loosely speaking, curvature can be inferred by the local behavior of geodesics – geodesics converge in regions of positive curvature and diverge in regions of negative curvature. On Riemannian manifolds, Ricci curvature is intimately related to the spread of geodesics emanating from the same point [20].

While there are many ways to characterize the local behavior of Ricci curvature, we focus on Ollivier’s characterization that is relevant for our purposes: namely that in regions of positive (resp., negative) Ricci curvature, geodesic balls (on average) are closer (resp., farther) than their centers [20]. (A “geodesic ball” of radius *ϵ* centered at a given point *p* is defined as the image under the exponential map of the ball of radius *ϵ* on the tangent space at *p*). This is in contrast to Euclidean space where the distances between geodesic balls and their centers are the same. Ollivier’s characterization generalizes this notion of Ricci curvature applicable to graphs by replacing the geodesic balls with probability measures *μ*_*j*_ [6]. In the Euclidean case, one may think of this as replacing points (delta functions), by small Gaussian balls (“fuzzified points”). The transportation distance between measures *μ*_*j*_ and *μ*_*k*_, prescribed by the Wasserstein distance *W*_1_, is used in lieu of the average distance between geodesic balls. The Wasserstein distance accounts for the geometry of the space and the distance between distributions associated with two nodes is related to the overlap of their neighborhoods. The rigorous mathematical details will be given now.

### 2.1 Wasserstein distance

The Wasserstein distance is a particular instance of the *optimal mass transport* (OMT) problem. It is a natural candidate for comparing probability measures because it accounts for both the shape of the distributions (i.e., weighted values) and the distance on the underlying space. The OMT problem, originated by Gaspard Monge [21], seeks the optimal way to redistribute mass with minimal transportation cost. Leonid Kantorovich reformulated and relaxed the problem in the context of resource allocation [22]; for more details, see [23, 24, 25]. We consider the following discrete formulation. Since we will be applying the theory to weighted graphs, this will be sufficient.

Accordingly, let 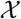 denote a metric measure space equipped with distance *d*(*·, ·*). Given two (discrete) probability measures *μ*_0_ and *μ*_1_ on 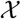, the ***Wasserstein distanc*** *W*_1_ between *μ*_0_ and *μ*_1_ is defined as

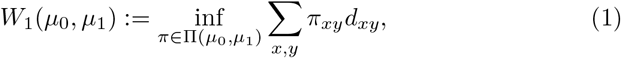

where ∏(*μ*_0_, *μ*_1_) is the set of joint probabilities on 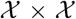 with marginals *μ*_0_ and *μ*_1_. Here, *π*_*xy*_ may be interpreted as the amount of mass moved from *x* to *y* and the cost of transporting a unit of mass is taken to be the distance travelled (i.e., *d*). Thus, the Wasserstein distance (1) gives the minimal net cost of transporting mass distributed by *μ*_0_ to match the distribution of *μ*_1_. The OMT problem therefore seeks the optimal *transference plan* ∈ ∏(*μ*_0_, *μ*_1_) found to be the infimal argument for which the Wasserstein distance is realized.

### 2.2 Curvature

The interplay between Ollivier-Ricci curvature, network entropy and functional robustness is linked by optimal mass transport (OMT), and is rich in theory. We outline this now, beginning with the Ollivier-Ricci curvature [6].

Based on the work of von Renesse and Sturm [16], Ollivier extended the notion of Ricci curvature, defined on a Riemannian manifold, to discrete metric measure spaces [6]. Specifically, let 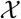 be a metric measure space equipped with a distance *d* such that for each 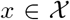, one is given a probability measure *μ*_*x*_. The probability measure *μ*_*x*_ can be thought of as *fuzzifying* or *blurring* the point *x*. For two points 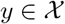, ***Ollivier-Ricci curvature*** is defined as

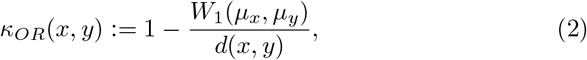

where *W*_1_ is the *Wasserstein distance*.

### 2.3 Curvature on graphs

For our purposes, the metric measure space is taken to be a weighted graph *G* = (*V, E*) with nodes (vertices) *V* and edges *E*. *G* is assumed to be a *simple, connected* and *undirected* graph. Instead of points *x* in a metric space, we now consider nodes *x*_*j*_ ∈ *V*, denoted simply by its subscript *j*. In this work, the graph is constructed as follows. Each node *j ∈ V* represents a gene; hereafter node and gene are used interchangeably. Edges *e* = (*j, k*) ∈ *E* define known interactions between genes (nodes) at the protein level (here given by HPRD) and *j ~ k* denotes that *k* is a neighbor of *j*. We then incorporate copy number (CN) values as nodal weights, denoted *w*_*j*_. Note that for *j ∈ V*, we take *w*_*j*_ = (*CN*)_*j*_ + 1; the affine translation is used to ensure all weights are positive.

We treat the weighted graph as a Markov chain. In this context, the probability measure *μ*_*j*_ attached to node *j ∈ V* can be thought of as the probability of a 1-step random walk starting from node *j*. The 1-step transition probability *p*_*jk*_ of going from *j* to *k* is expressed by the *principle of mass action* [26].

According to this principle, if there is a known connection between gene *j* and gene *k* (i.e., (*j, k*) ∈ *E*), then the probability that they interact is proportional to the product of their CN values:

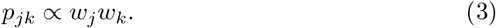

Normalizing the mass action over all possible edges to ensure that *p*_*jk*_ is a probability, i.e., ∑_*j*~*k*_ *p*_*jk*_ = 1, we define the transition probabilities *p*_*jk*_ of the stochastic matrix *P* = [*p*_*ij*_] associated with the Markov chain as follows:

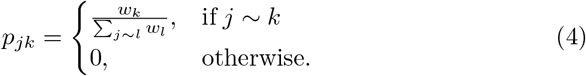

Accordingly, for each gene *j*, we associate a probability measure *μ*_*j*_ defined on the node set *V* with *n* associated nodes

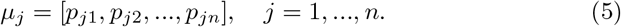

Alternatively, *μ*_*j*_ can be thought of as *fuzzifying* the node *j* over its 1-step neighborhood.

#### 2.3.1 Graph distance

We have now specified the points (*x*) and measures (*μ*_*x*_) needed to compute OR curvature in Eq. (2) on a graph. All that remains is the distance *d*(*x, y*). In lieu of the commonly used *hop distance*, i.e., the distance between two nodes *j, k* ∈ *V* that is defined as the shortest path length over all paths connecting *j* and *k*, we take the corresponding graph distance *d*_*jk*_ to be the ***weighted hop distance*** (whop).

More precisely, for fixed nodes *j* and *k*, let *P*^*jk*^ denote a path connecting them. Let 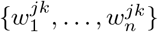 be the set of all the associated edge weights. Then we set

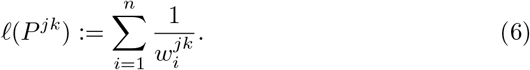

Denoting by 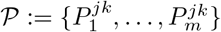, the set of all possible paths connecting *j* and *k*, we define the ***weighted hop distance (whop)*** between *j* and *k* to be:

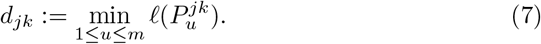

Note that the edge weights *w*_*uv*_ for all edges *e* = (*u, v*) ∈ *E* are constructed as

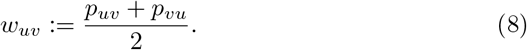

This formulation was chosen so the distance between two nodes is inversely related to the probability of their interaction. Thus, the higher (resp., lower) the probability of two nodes interacting, the smaller (resp., larger) the distance between them should be. The average is taken merely so the distance is symmetric, i.e., *d*_*jk*_ = *d*_*kj*_.

Using Zachary’s Karate Club graph [27] as an example, the resulting whop distance for all edges is shown in Figure 2. A more detailed comparison between the hop and whop distances, illustrated by heat maps of the corresponding distance matrices of all node pairs in the network, is shown in Figure 3.

**Figure 2:**
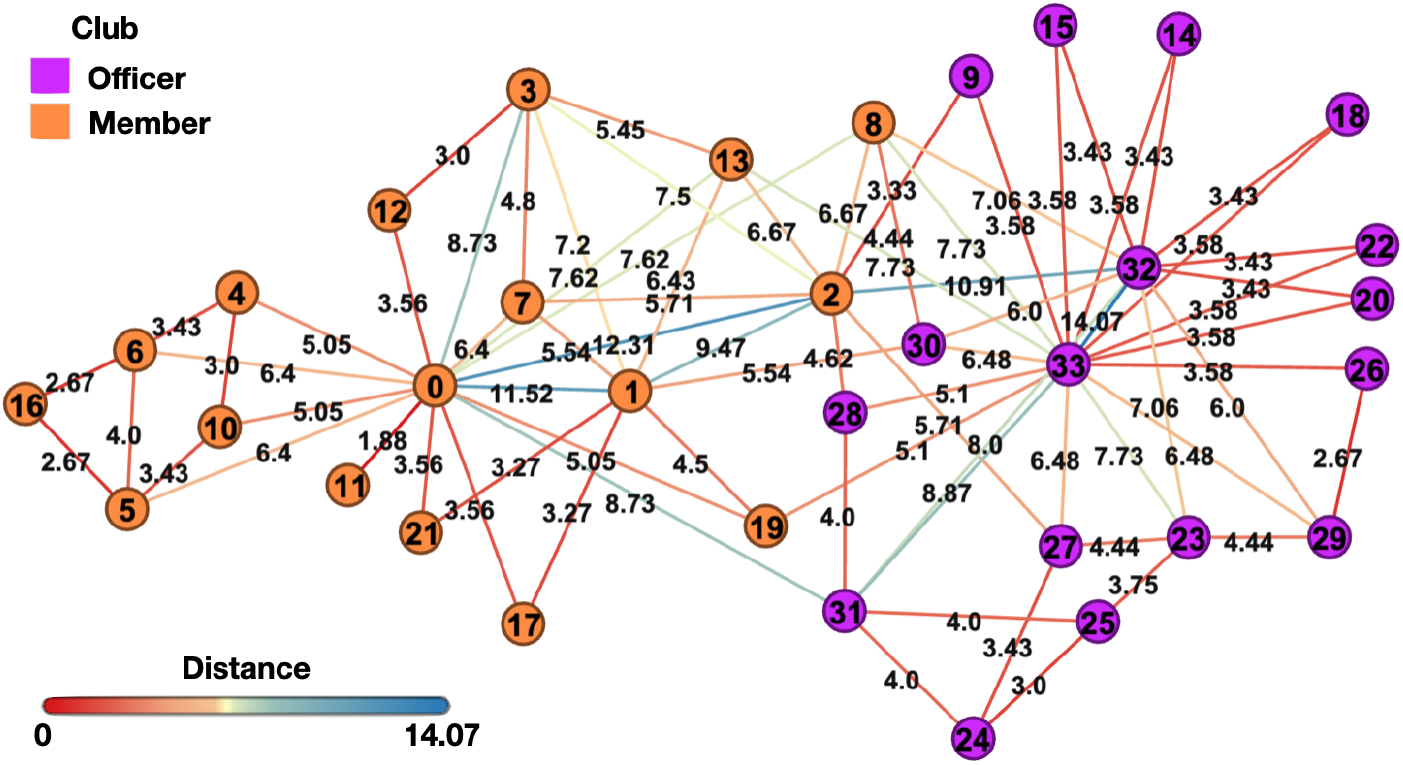
Weighted hop distances are shown for every edge in Zachary’s Karate Club Graph [27] with all node weight values initialized equal to 1. The node color indicates if the corresponding person is a club officer (purple) or member (orange). The distance between edge-adjacent nodes is shown at the edge midpoint.

**Figure 3:**
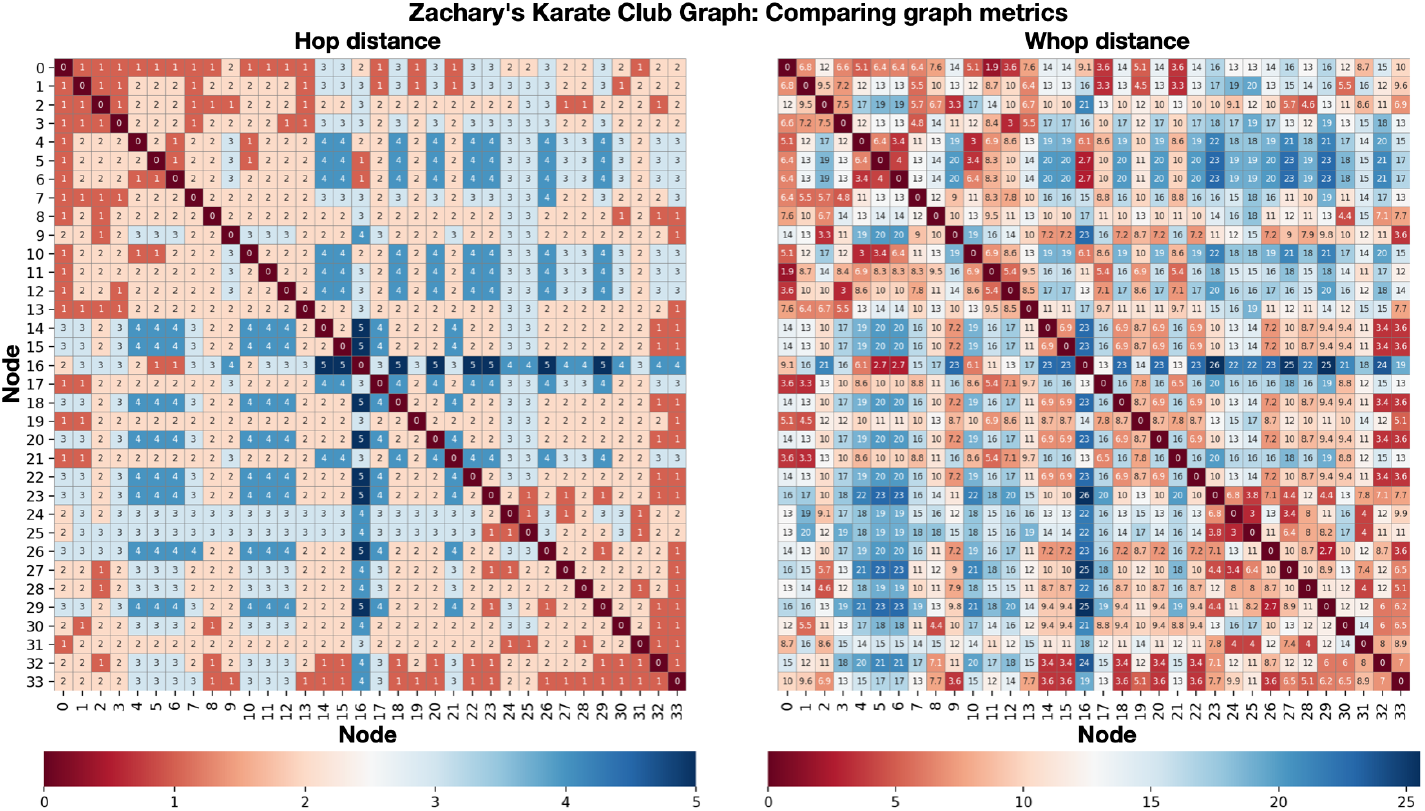
Comparing graph metrics. The distances between every two nodes in Zachary’s Karate Club Graph [27], with all node weight values initialized equal to 1, are shown using (left) the hop distance and (right) weighted hop distance.

#### 2.3.2 Edge curvature

With the choice of graph distance in Eq. (7), the OR curvature in Eq. (2) can now be computed between any two nodes in the graph. Due to the large nature of the graphs of interest, we constrain the curvature computation to edges. Notice, from the curvature definition in Eq. (2), the ratio 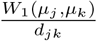 relates the transport cost of moving the distribution (i.e., *fuzzy* ball) associated with *j* to *k* to the ground distance. Informally, the more the neighborhoods of two nodes overlap, the lower the transportation cost between them and thus the higher the curvature associated with the edge. As such, curvature informs on the local functional relationship between neighborhoods.

#### 2.3.3 Scalar and total curvature on graphs

In order to obtain a node-level measure of curvature, we consider a contraction of the edge curvatures, analogous to *scalar curvature* defined on points of a manifold in Riemannian geometry [20]. In this work, we define the (nodal) ***scalar curvature*** of gene *j* to be the weighted sum of the curvatures on all edges incident to *j*:

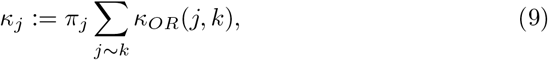

where the weight *π*_*j*_ is the *j*^*th*^ component of the stationary distribution *π* associated with the Markov chain *P* [26]:

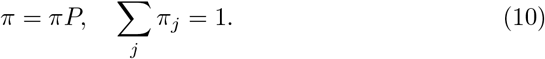

The stationary distribution in this setting (connected graph) is also the limiting distribution of the Markov chain, known as the *stationary* or *equilibrium* distribution. Thus, the quantity *π*_*j*_ describes the relative importance of node *j* with respect to all other nodes. We therefore scale the nodal curvature by its component in the stationary distribution in order to correct for nodal bias. Furthermore, the stationary distribution has a closed form that may be easily computed as follows:

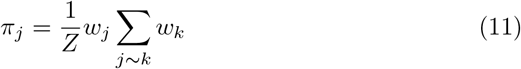

where *Z* is the normalization factor. We note that unweighted and alternative weightings have been proposed [28, 29].

Lastly, we define the ***total curvature*** *κ*_*G*_ of a network to be the net scalar curvature, summed over all nodes in the graph

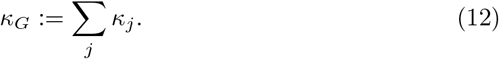

### 2.4 Curvature and robustness

Sturm [16], Lott and Villani [15] related a lower bound on the Ricci curvature of a smooth Riemannian manifold to the entropy of densities along a constant-speed geodesic with the use of the Wasserstein distance. This laid the groundwork for the connection between curvature, entropy, and the Wasserstein metric, and led to the remarkable observation that changes in Ricci curvature Δ*κ*_*Ric*_ are positively correlated with changes in (Boltzmann) entropy Δ*S*:

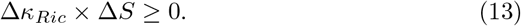

The positive correlation between changes in curvature Δ*κ*_*Ric*_ and changes in robustness Δ*R*:

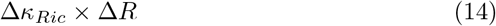

is realized by Eq. (13) and the fluctuation theorem [30] from large deviations theory indicates that changes in entropy are positively correlated with changes in robustness Δ*R* :

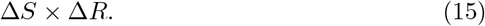

Here, *robustness* refers to the ability of a system to recover or maintain its ability to function after it is perturbed in some way (e.g., stress signal).

Curvature is a particularly attractive method for analyzing key nodes and interactions in large complex PPI networks primarily due to its intimate connection to robustness. This connection is linked by entropy as shown in Eqs. (13, 15), bridging this geometric analysis to an interesting perspective on the relationship between the topological and functional properties of the weighted network. With this notion of the change in curvature as a proxy for the more qualitative notion of functional robustness, we rank genes according to the change in curvature with respect to the topology and between sub-groups identified; see the following Results Section.

### 2.5 Data description and processing

In this section, we outline the data description and processing that we used in our HGSOC analysis. Further details about the data may be found in [12].

First of all, TMB was calculated by dividing the number of non-synonymous mutations by the total size of the capture panel in megabases. Secondly, based on the CNAs by FACETS, the fraction of genome altered (FGA) was defined as the cumulative length of segments with log 2 or linear CNA value larger than 0.2 divided by the cumulative length of all segments measured. Large-scale state transition (LST) scores, defined as a chromosomal breakpoint resulting in allelic imbalance between adjacent regions of at least 10Mb, were determined, and a cut-off ≥ 15 was employed for LST-high cases.

Next, regarding the data characteristics, we used DNA gene CNA data from a subset of 69 women with recurrent OC who received immunotherapy from a previously published series [12]. The subtypes of ovarian cancer are in fact quite different diseases, originating in different cell types and being caused by distinct mutations with diverse outcomes, and should therefore be analyzed separately [19]. Accordingly, we restrict our re-analysis to a subset of samples (*n* = 49) with HGSOC, which is the most common and lethal subtype. Four HGSOC patients had two samples, and the replicate samples were removed from the analysis. This resulted in a total of 45 tumor samples, 32 of which were metastases and 13 represented primary (adnexal) tumors, with 22 and 10 deaths in each group, respectively, at the time the study group was analyzed. This forms a homogeneous group of cancers (Table 1). Tumor and normal samples from the 45 patients were profiled utilizing the FDA-cleared Memorial Sloan Kettering Integrated Mutation Profiling of Actionable Cancer Targets (MSK-IMPACT) sequencing assay, their mean age was 58 years, and mean TMB was 5.9. Patient selection and clinical characteristics are displayed in Figure 4 and in Tables 1,2, respectively.

**Table 1:**
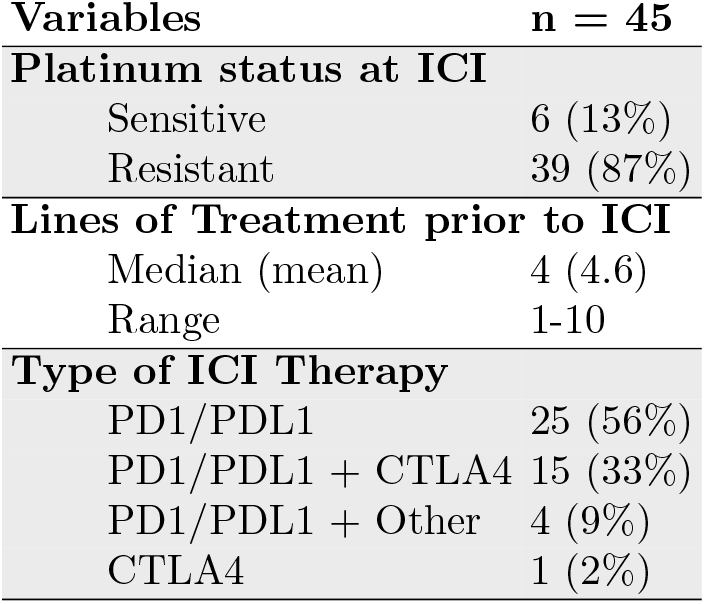
Clinical characteristics of patients with recurrent high grade serous ovarian cancer administered immune checkpoint inhibitor (ICI) therapy.

**Figure 4:**
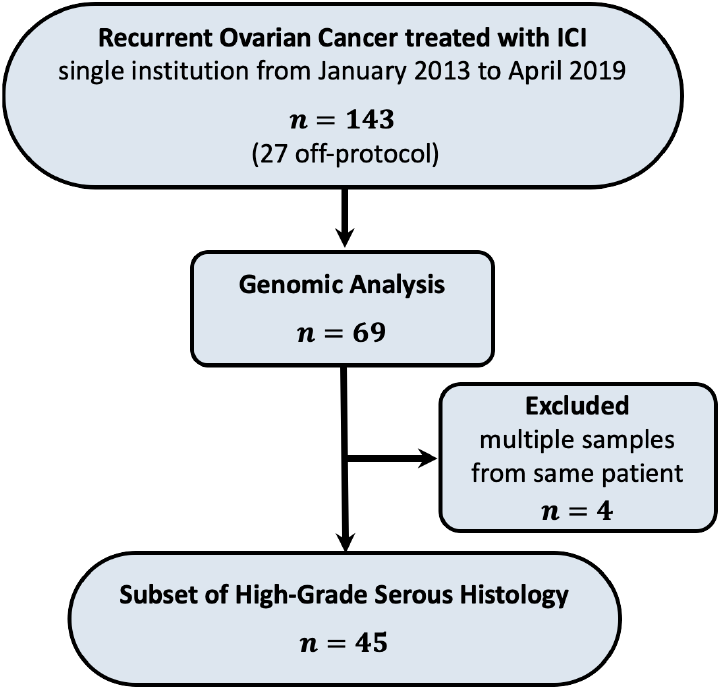
Patient selection

**Table 2:** HGS patient characteristics. Abbreviations: SD, standard deviation; IQR, interquartile range. *P*-values were obtained using two-sided Wilcoxon-Rank Sum test for continuous variables and Fisher-exact test for categorical variables.

| Characteristic | All Patients<br>(n=45) | Low curvature<br>(n=12) | High curvature<br>(n=33) | *p* |
| --- | --- | --- | --- | --- |
| Age at diagnosis (years) |  |  |  | 0.062 |
| Mean ± SD | 58.0 ± 9.3 | 62.3 ± 7.5 | 56.4 ± 9.4 | |
| Range | 27.0–75.0 | 49.0–75.0 | 27.0–75.0 |  |
| Median (IQR) | 58.0 (52.0–64.0) | 64.0 (58.8–65.5) | 55.0 (51.0–61.0) |  |
| Age at start of ICI (years) |  |  |  | 0.023 |
| Mean ± SD | 62.1 ± 8.7 | 67.1 ± 6.9 | 60.3 ± 8.7 | |
| Range | 37.0–78.0 | 55.0–78.0 | 37.0–77.0 |  |
| Median (IQR) | 62.0 (56.0–69.0) | 67.0 (63.3–70.3) | 59.0 (55.0–66.0) |  |
| Stage at diagnosis |  |  |  | 0.502 |
| III | 25 | 8 | 17 |  |
| IV | 20 | 4 | 16 |  |
| Time from diagnosis to start of ICI (months) |  |  |  | 0.581 |
| Mean ± SD | 50.9 ± 35.9 | 58.8 ± 43.1 | 48.1 ± 33.2 | |
| Range | 5.3–166.0 | 17.4–166.0 | 5.3–123.1 |  |
| Median (IQR) | 49.4 (23.3–61.7) | 44.5 (33.4–61.8) | 49.4 (18.5–61.7) |  |
| Duration of ICI (weeks) |  |  |  | 0.807 |
| Mean ± SD | 20.2 ± 23.6 | 14.3 ± 8.6 | 22.3 ± 27.0 | |
| Range | 0.1–143.0 | 0.7–28.3 | 0.1–143.0 |  |
| Median (IQR) | 12.3 (7.7–23.1) | 13.6 (7.8–20.3) | 12.1 (7.7–23.1) |  |
| Overall survival (months) |  |  |  | 0.007 |
| Mean ± SD | 16.7 ± 11.7 | 8.8 ± 7.2 | 19.6 ± 11.7 | |
| Range | 0.4–44.8 | 0.4–27.4 | 0.4–44.8 |  |
| Median (IQR) | 15.3 (6.5–24.6) | 7.4 (4.9–10.9) | 20.3 (11.1–26.0) |  |
| Sample type |  |  |  | 0.010 |
| Metastasis | 32.0 | 12 | 20 |  |
| Primary | 13.0 | 0 | 13 |  |
| Status at last follow-up |  |  |  | 0.134 |
| Alive | 13 | 1 | 12 |  |
| Dead | 32 | 11 | 21 |  |
| TMB |  |  |  | 0.959 |
| Mean ± SD | 3.7 ± 2.3 | 3.5 ± 1.9 | 3.8 ± 2.5 | |
| Range | 1.0–9.7 | 1.1–6.7 | 1.0–9.7 |  |
| Median (IQR) | 3.3 (2.0–4.4) | 2.6 (2.0–5.3) | 3.3 (2.0–4.4) |  |
| FGA |  |  |  | 0.005 |
| Mean ± SD | 0.4 ± 0.2 | 0.3 ± 0.2 | 0.5 ± 0.2 | |
| Range | 0.005–0.871 | 0.005–0.629 | 0.092–0.871 |  |
| Median (IQR) | 0.4 (0.3–0.6) | 0.2 (0.1–0.4) | 0.5 (0.4–0.6) |  |
| LST |  |  |  | 0.024 |
| Mean ± SD | 25.0 ± 10.3 | 19.3 ± 9.1 | 27.1 ± 10.0 | |
| Range | 2.0–51.0 | 2.0–32.0 | 2.0–51.0 |  |
| Median (IQR) | 25.0 (20.0–29.0) | 22.0 (13.5–25.8) | 27.0 (22.0–34.0) |  |

CN segments were mapped to individual genes according to GRCh37 and for each sample, each gene was assigned the maximum CN value of all segments that mapped to it. After removing all genes with missing data and all genes not in the HPRD network, we extracted the set of genes comprising the largest connected network (Supplementary Figure S7). This resulted in a CNA data matrix of size 3,489 (genes) × 45 (samples).

The network topology was constructed as follows. Edges between genes were defined by the PPI obtained from HPRD [1, 2]. The network topology was then taken to be the largest connected component in the HPRD network restricted to the set of genes in our data set. This resulted in a network with 9,710 edges and 3,489 nodes with an average degree of 5.57. The rationale is that the established interactions between gene products serve as a viable proxy for the functional connectivity at the gene level.

Subject specific networks were created by assigning nodal weights *w*_*j*_ prescribed by the CN value. Specifically, the CN data took on discrete integer values in the range [0, 38]. In order to ensure all weights were positive, we used the translation *w*_*j*_ = *x*_*j*_ +1 where *x*_*j*_ is the CN value for gene *j*. For each subject, Markov chains were computed as defined in Eq. (4) followed by the associated stationary distribution in Eq. (11). Next, Ollivier-Ricci curvature using Eq. (2) was computed on each edge in the fixed network, scalar curvature defined in Eq. (9) was subsequently computed for each node and lastly, total curvature using Eq. (12) was computed for the network. A critical aspect of the curvature analysis is that it provides a *relative* quantity and it is the *change* in curvature that is of interest, indicative of changes in the network’s capacity for communication. Thus, we would expect that patients whose samples have a lower total curvature (i.e., a relative net decrease in capacity) would be associated with a poorer prognosis than those with higher total curvature values.

## 3 Results

### 3.1 Survival analysis

The prognostic value of the total curvature *κ*_*G*_ in Eq. (12) and standard genomic parameters including TMB, FGA and LST (representing homologous recombination deficiency [HRD] status) were assessed with respect to the HGS cohort (*n* = 45). For each parameter (TMB, LST, FGA, *κ*_*G*_), the cohort was stratified into two groups according to the 25th percentile (low vs. high) of individual values. The cutoff was selected based on the location where the curve fitted to the sorted total curvature values starts slowly incrementing and is approximately linear (Supplementary Figure S3). An alternative cut point using maximally selected log-rank statistic [31, 32] was assessed as well and resulted in a comparable split (Supplementary Figure S4). However, a larger cohort is needed for further validation. The effectiveness of each parameter in terms of OS was evaluated using the Kaplan-Meier (KM) analysis [33].

OS was defined from the start of immunotherapy treatment until either death or last follow-up [12]. Survival curves for each parameter were plotted according to the KM estimator, shown in Figure 5 along with the corresponding log-rank p-values (total curvature: *p* = 0.00047; TMB: *p* = 0.03153; LST: *p* = 0.42865; FGA: *p* = 0.19568). While both TMB and total curvature *κ*_*G*_ were found to be significant factors in predicting patient survival, the p-value for total curvature was almost 2 orders of magnitude smaller as compared to TMB, whose p-value was just marginally significant. The effective prognostic predictive power of the total curvature, particularly in comparison to the genomic parameters, is one of the major contributions of this work. See Supplementary Information for validation.

**Figure 5:**
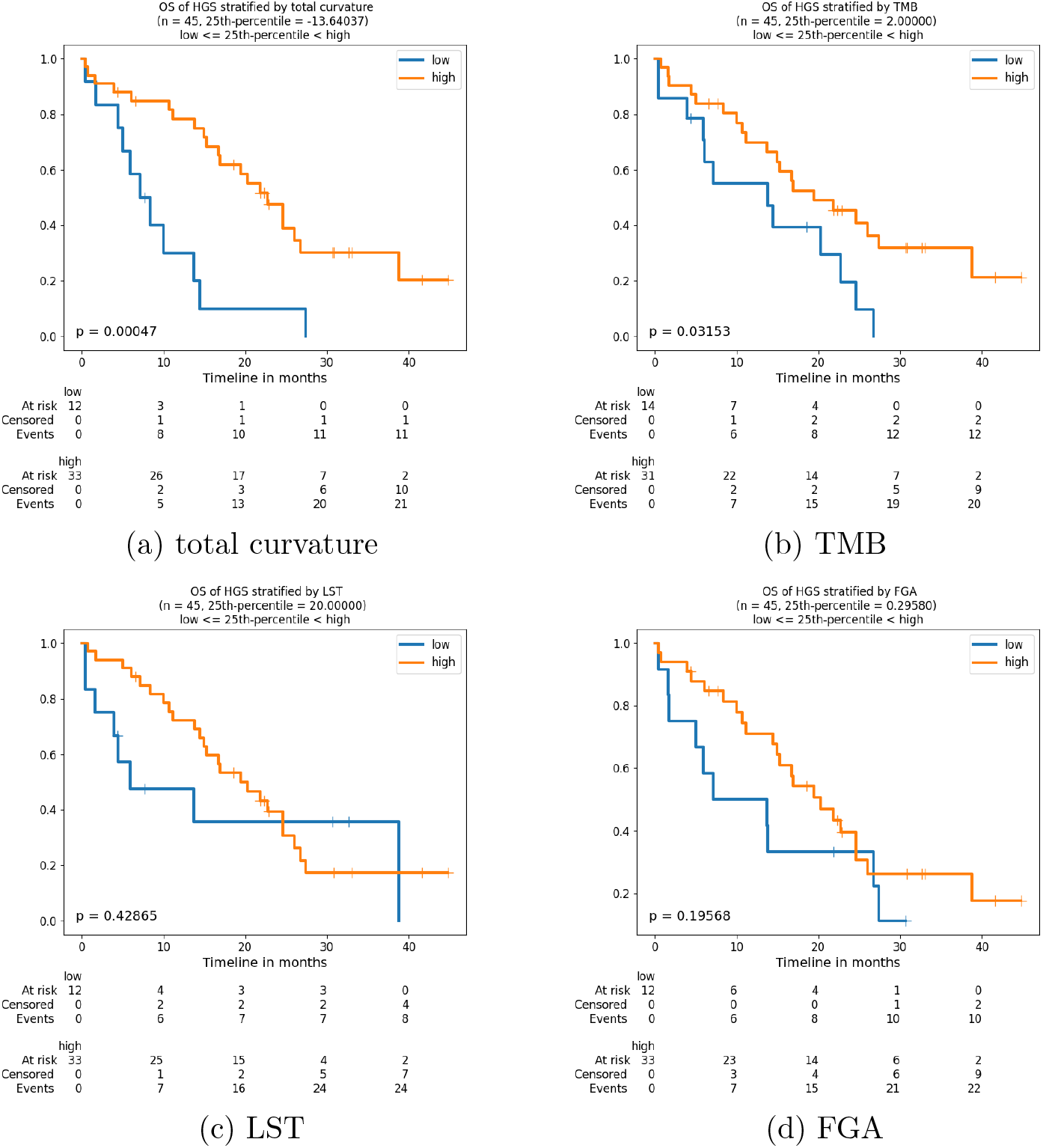
Survival curves for HGS samples (*n* = 45) stratified low and high groups by the 25th percentile of total curvature and genomic parameters. P-values were derived from the log-rank test.

In order to assess that the prediction is not independent of receiving immunotherapy treatment, we repeated the curvature and survival analysis pipeline on IMPACT data from HGSOC samples that did not receive ICIs. It is interesting to note that total curvature was not predictive of survival in this setting (Supplementary Figure S5), highlighting that our findings may be immunotherapy-specific. However, it is also important to point out that OS was defined from the time of diagnosis for the analysis of this dataset, whereas in the analysis of 45 HGSOC patients treated with ICIs, OS was defined from the start date of immunotherapy, and all 45 patients had recurrent tumors with a substantial time gap between the time of first diagnosis and the start date of immunotherapy.

### 3.2 Functional biomarkers

Genes that exhibit large changes in scalar curvature are identified as the genes that potentially play a key role in altering the network robustness (i.e., functional connectivity). This requires a reference for comparison, typically using data collected at a reference time (e.g., after immunotherapy treatment) or data collected from a reference sample (e.g., normal tissue). Often no such reference data are available, as was the case here where CNA data from only one time point were provided. Considering the distinction in survival curves obtained via curvature, we therefore used the high and low risk groups (as previously defined by the 25th percentile of the total curvature and dichotomized into low and high curvature groups, respectively) for points of comparison. Genes were ranked by the difference in average scalar curvature between the high and low risk groups (Δ*κ*_*risk*_). The change in curvature measures the relative gene implication in the stabilization (or destabilization) of local network robustness driving changes in feedback connectivity pertaining to survival. Since both increased and decreased functionality is of interest, the top 50 ranked genes that exhibited the largest positive (Δ*κ*_*risk*_ > 0) and largest negative (Δ*κ*_*risk*_ < 0) change in curvature, yielding 100 candidate genes associated with risk, are listed in Table 3).

**Table 3:** Changes in average scalar curvature based on risk (high vs low). Top 50 genes ranked by positive (Δ*κ*_*risk*_ > 0) and negative (Δ*κ*_*risk*_ < 0) difference in average scalar curvature between low risk (*n* = 33) and high risk (*n* = 12)

| rank | gene | $\Delta\kappa_{risk} > 0$ | gene | $\Delta\kappa_{risk} < 0$ |
| --- | --- | --- | --- | --- |
| 0 | TP53 | 0.208647 | CREBBP | -0.064223 |
| 1 | ATXN1 | 0.102823 | SHC1 | -0.031456 |
| 2 | EP300 | 0.044184 | PTK2 | -0.026202 |
| 3 | SMAD2 | 0.042756 | AR | -0.025316 |
| 4 | PIK3R1 | 0.037015 | MYC | -0.022608 |
| 5 | SRC | 0.033112 | JUN | -0.019546 |
| 6 | SMAD4 | 0.032177 | LYN | -0.011148 |
| 7 | RB1 | 0.031043 | YWHAQ | -0.010984 |
| 8 | ESR1 | 0.027914 | GSK3B | -0.009017 |
| 9 | PRKCA | 0.027253 | STAT1 | -0.008248 |
| 10 | CTNNB1 | 0.025121 | CDK5 | -0.007480 |
| 11 | GRB2 | 0.016848 | FN1 | -0.006947 |
| 12 | YWHAE | 0.015125 | COPS6 | -0.006251 |
| 13 | DLG4 | 0.014966 | SMAD3 | -0.006133 |
| 14 | PRKCD | 0.014742 | PAK1 | -0.006091 |
| 15 | ACTB | 0.013456 | MYOC | -0.005464 |
| 16 | EWSR1 | 0.012300 | SMURF1 | -0.005438 |
| 17 | TGFBR1 | 0.010799 | SUMO1 | -0.004455 |
| 18 | RAC1 | 0.008937 | PARP1 | -0.004274 |
| 19 | PLCG1 | 0.008423 | CRMP1 | -0.004271 |
| 20 | CHD3 | 0.007997 | HSF1 | -0.004155 |
| 21 | DVL2 | 0.007476 | HIPK2 | -0.004038 |
| 22 | BCL2 | 0.007009 | CDC42 | -0.004017 |
| 23 | RANBP9 | 0.006879 | POU2F1 | -0.003838 |
| 24 | MAPK1 | 0.006630 | ACVR1 | -0.003651 |
| 25 | POLR2A | 0.006468 | HTT | -0.003537 |
| 26 | CRK | 0.006375 | JAK1 | -0.003520 |
| 27 | APP | 0.006256 | PDPK1 | -0.003497 |
| 28 | PCNA | 0.005935 | PIK3R2 | -0.003423 |
| 29 | COIL | 0.005350 | FGFR1 | -0.003352 |
| 30 | MAPK14 | 0.005097 | CDKN1A | -0.003205 |
| 31 | NR3C1 | 0.004981 | MAGEA11 | -0.003165 |
| 32 | AKT1 | 0.004925 | GNAI1 | -0.003125 |
| 33 | EGFR | 0.004918 | PRKCE | -0.003090 |
| 34 | RHOA | 0.004635 | XPO1 | -0.002919 |
| 35 | RAF1 | 0.004159 | BTK | -0.002855 |
| 36 | SMAD7 | 0.004071 | MUC1 | -0.002814 |
| 37 | NCOR1 | 0.004038 | EIF2AK2 | -0.002807 |
| 38 | RASA1 | 0.003998 | CASP8 | -0.002758 |
| 39 | FXR2 | 0.003879 | CSNK2A2 | -0.002717 |
| 40 | RPA1 | 0.003560 | MDM2 | -0.002710 |
| 41 | HRAS | 0.003525 | NTRK1 | -0.002636 |
| 42 | UBB | 0.003302 | ADAM15 | -0.002541 |
| 43 | BRCA1 | 0.003292 | FASLG | -0.002522 |
| 44 | SUMO4 | 0.003283 | VIM | -0.002436 |
| 45 | ARRB2 | 0.003248 | CD247 | -0.002372 |
| 46 | XRCC6 | 0.003065 | AXIN1 | -0.002333 |
| 47 | HGS | 0.003025 | SMARCA4 | -0.002256 |
| 48 | HDAC3 | 0.002965 | SNAPIN | -0.002246 |
| 49 | HSP90AA1 | 0.002924 | PPP2R5A | -0.002187 |

Similarly, we investigated the top genes ranked by the difference in average scalar curvature between sub-groups based on available clinical data as an exploratory analysis. Of ancillary interest were the top ranked candidate driver genes that demonstrate functional network response to ICI and their association to survival as exhibited by disparities in network robustness measured between those who were alive or deceased at last follow-up (Δ*κ*_*OS*_; Supplementary Table S1) and predominant changes in functional connectivity due to DNA level dysregulation that occurs between primary and metastatic tumors (Δ*κ*_*PM*_; Supplementary Table S2).

Lastly, we used the network topology itself as a frame of reference. Treating the fixed network topology as an unweighted graph (i.e., all node weights are uniformly set to 1), we computed the scalar curvature on this reference topology network in the same manner as detailed above. This provides a measure of discordance in functional connectivity between the HGSOC network and its underlying topological structure (Δ*κ*_*ref*_; Supplementary Table S3). It is interesting to note that in all of the comparisons TP53 appeared at the top of all positive changes in curvature indicating its functional centrality in HGSOC.

Substantial overlap in the top 50 (positive and negative) ranked genes was noted from all of the comparisons performed, resulting in 171 unique genes listed in Supplementary Table S4 (Supplementary Figures S8,S9). The choice of selecting the top 50 genes was largely arbitrary with the following rationale. The assertion that critical genes may be identified as those exhibiting larger changes in curvature is supported by the theory, but curvature is a continuous variable with no obvious cutoff. Since there is also an exploratory component to this analysis, we opted for a cutoff that would yield a manageable set of genes that reasonably included the key influential players. Out of 3,489 genes in the network, this resulted in 50 (positive and negative) candidate genes. See Supplementary Figure S6 for a further *s* ub-curvature analysis on the association between the highlighted candidate genes and survival.

### 3.3 Relationship between total curvature and genomic features

Lastly, we explored the relationship between total curvature and genomic features (TMB, FGA, LST). Linear regression analysis with two-sided Wald test and Pearson correlation (*r*) analysis were used to assess the correlation between total curvature and each of the clinical features (TMB: *p* = 0.9674; FGA: *p* = 0.0060; LST: *p* = 0.0868). This analysis suggests that total curvature is significantly correlated with FGA. This result is not entirely surprising considering that FGA is a surrogate measure of CN changes and the curvature measures dysregulation of the CN-weighted network. However, total curvature yields high and low risk groups with a significant difference in survival whereas FGA does not. The difference is that total curvature accounts for an extra level of information, namely the connectivity, that is not evident from CNAs alone. We believe this is compelling evidence that network dysregulation, as measured by curvature, has the potential to provide critical insight for analyzing immune response. More samples are needed to verify this result but it is interesting to note that further investigation into FGA as a potential biomarker for survival in HGSOC has been proposed [12]. Linear regression plots on the HGS cohort (n=45) are shown in Figure 6.

**Figure 6:**
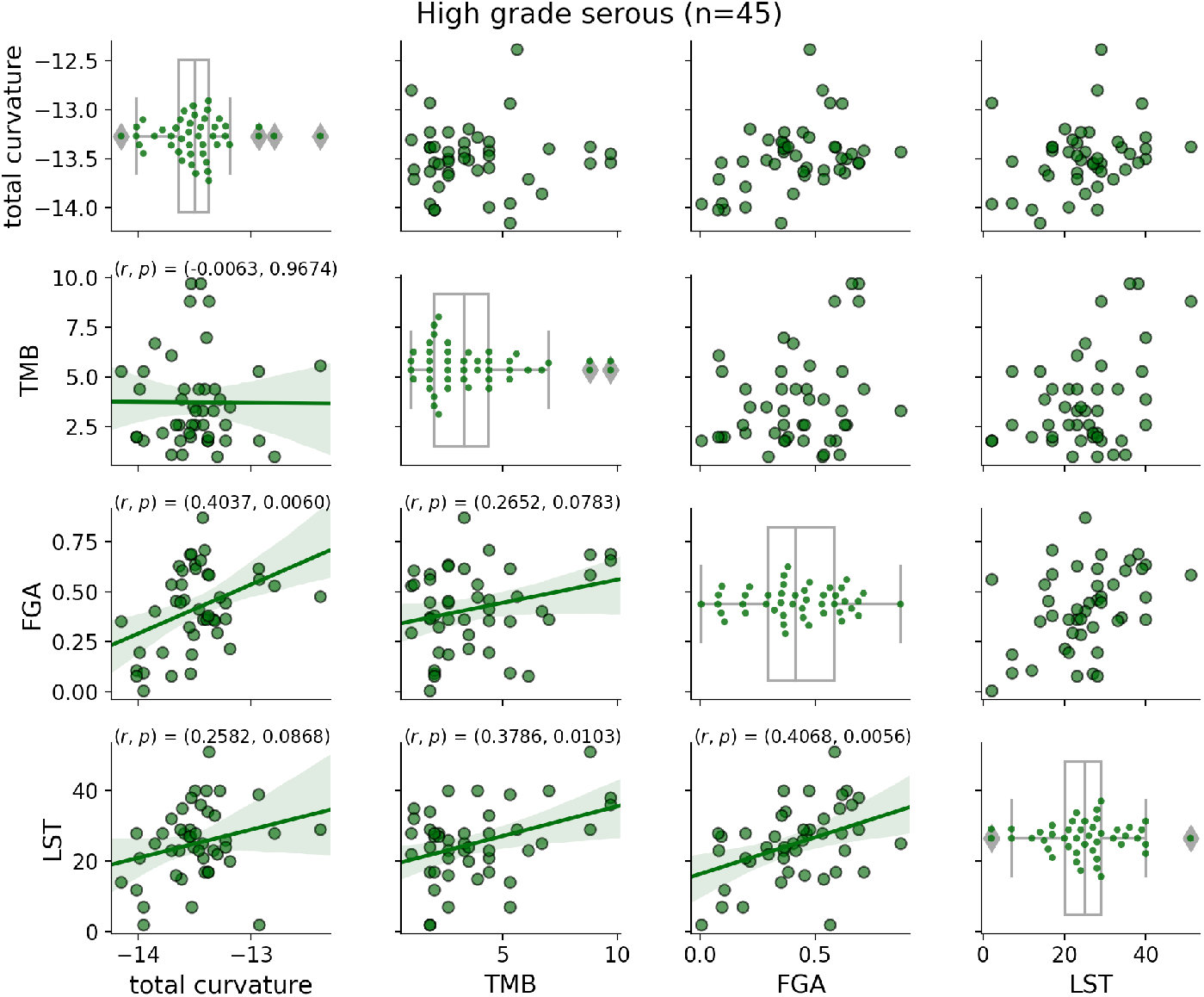
Linear regression of total curvature onto clinical parameters using HGS samples (n=45). The lower triangle includes the Pearson correlation (*r*) and two-sided p-value for the hypothesis test with *H*_0_ : the slope is zero, using Wald test with t-distribution of the test statistic and 95% confidence interval.

## 4 Discussion

### 4.1 Biological/molecular relevance

Mutational profiles of HGSOCs are characterized by abnormal gene CNAs, which results in protein overexpression or underexpression [13]. The majority of these OCs are characterized by inactivating mutations or loss of TP53, leading to aneuploidy, resulting from loss of control of centrosome numbers [34], and selection for enhanced copy number and gene expression of selected genes controlling the cell cycle (Figure 7). These OCs commonly overexpress the cyclin E protein due to loss of p53 function, resulting in downregulation of p21 (the inhibitor of cyclin E-Cdk-4/6 activity), as well as amplification of cyclin E [13]. In addition, the serous OCs have one or more of the K-RAS, MYC, and AKT protein kinase genes overexpressed in the late G-1 phase of the cell cycle (see Figure 7). The K-RAS activity signals that the cell is stimulated by growth factors and should progress through the cell cycle, the MYC gene regulates the transcription of hundreds of genes for cell growth and division and the AKT gene promotes TORC-2 activity for entry into S-phase and stimulates AKT ki-nase to enhance the MDM-2 E3 ubiquitin ligase to increase the destruction of the p53 protein [35]. All of these driver gene products promote a constant over-expressed signal for cell cycle progression and division. The mutational profile of this cancer is copy number changes of genes and overexpression of selected gene products. For that reason, the methods developed here employ copy number values as the measurement for each node containing a gene in the signal transduction pathway and the resultant network that is employed to measure curvature.

**Figure 7:**
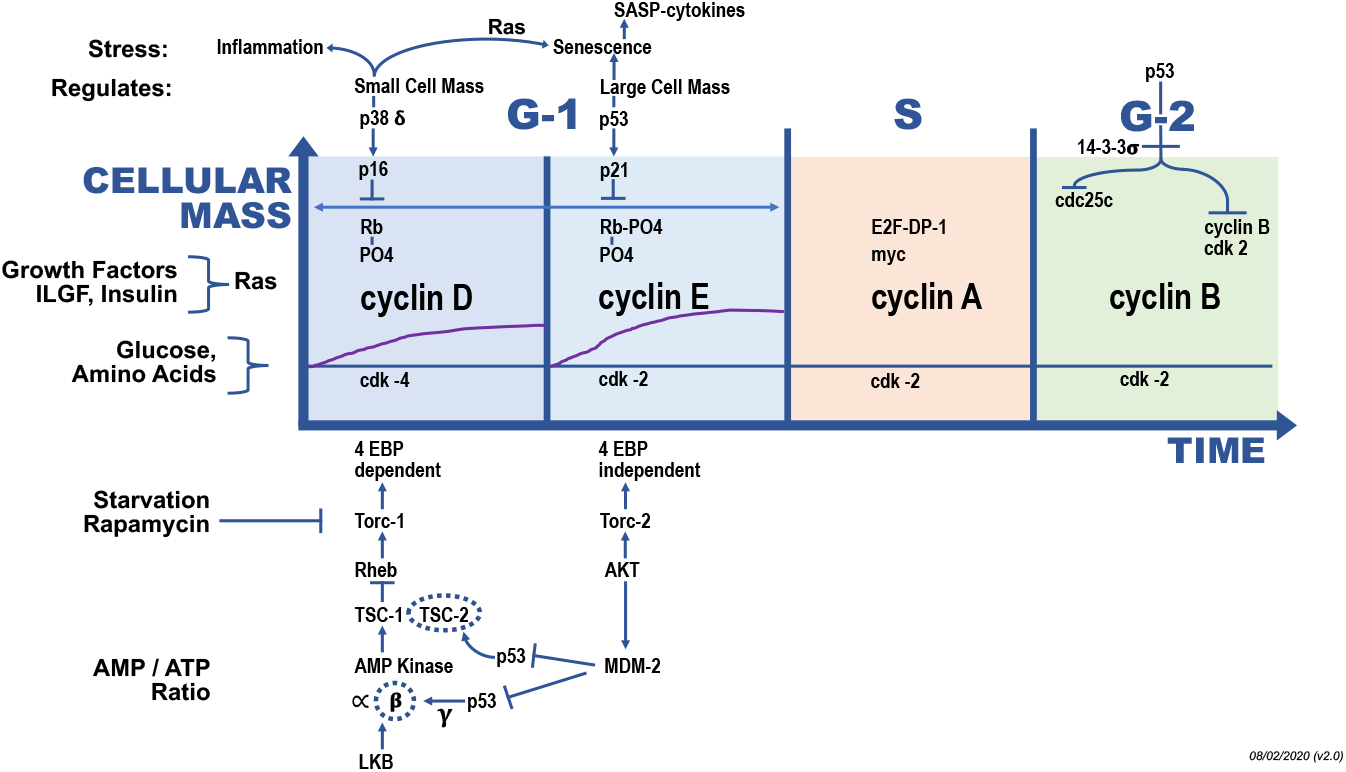
Genes involved in Serous Ovarian Cancer in the G-1 Phase of the Cell Cycle: The G-1 phase of the cell cycle can be divided up into cyclin D-cdk4/6 early events and cyclin E-cdk2 later events. The inhibitors of these protein kinase activities, p38 and p16 for cyclin D and p53 and p21 for cyclin E are shown above the cyclin D and E panels. The activating pathways for cyclin D (TORC-1) and cyclin E (TORC-2) are shown below these panels. The mutational loss of TP53 and the amplification of cyclin E results in the loss of control of cyclin E levels and the hyper-amplification of centrosome numbers destabilizing the copy number control of chromosome numbers (aneuploidy) and gene copy numbers. Serous ovarian cancers commonly have K-RAS, MYC and AKT genes or chromosome amplifications and overexpression. The p21 gene is not mutated suggesting that it has additional functions required elsewhere for viability or that additional functions of p53 must be lost for ovarian cancers. Every gene highlighted in this figure can be found genetically altered in a cancer of other tissue types.

This mutational profile of serous OC results in the loss of control for duplicating centrosomes, which sets up the polarity in a cell for the normal segregation of chromosomes. This is driven by the loss of function of p53 and the overexpression of cyclin E, which co-localizes with the centrosome, which duplicates abnormally producing three or more centrosomes [36]. In the extreme, this results in chromothripsis, where a chromosome fragments and some of the parts are reassembled in a random order. This can result in double minute chromosomes without a centromere for proper segregation and random partition of the double minutes and distribution of multiple copy numbers. Often the population of cells forms a distribution of copy numbers of a combination of genes, which are then selected for optimal fitness.

Biomarkers of response to immunotherapy in OC remain underdeveloped. Here, we characterized a cohort of HGSOC patients treated with immunotherapy for whom detailed treatment, genomic, and survival data were available. Our analysis indicates that employing the copy number of the relevant genes as a measurement for each node in a network provides the strongest predictive power for OS, when compared to prior examined parameters such as TMB, LST, and FGA (Figure 5). These results suggest that no one gene or even its alterations can predict responses to therapy. Rather it is the integration of the copy numbers of driver genes and the change of resultant networks formed by these genetic or epigenetic alterations that impacts immunological responsiveness of the tumor after checkpoint therapy. Employing the overexpression of the same set of genes and loss of p53 function in a mouse model of ovarian cancers treated with immunotherapy resulted in similar heterogeneous responses to checkpoint therapy and the beginnings of experimental tests of genes and products that could modify the results of the responses to cancer therapies [14]. This permits the pairing and testing of the type of modeling presented here along with prediction of genes with high curvature with experimental tests in a mouse model to improve the choice of therapies depending upon the genotypes of the tumors.

Interestingly, in non-small cell lung cancer a major tumor antigen, not genetically altered in sequence (not a neo-antigen), was found to be overexpressed in many different independent tumors [7, 8]. This suggests that in serous OCs, like non-small cell lung cancers, the higher concentration of a non-genetically altered tumor antigen was an important variable in responsiveness to checkpoint therapy. Similar conclusions were reached by the mathematical construct employed here and measured by both abundance and changes in a network architecture and quantitated by curvature of the edges of the network.

### 4.2 Conclusions

The marriage of mathematical models with experimental tests is one of the goals that will speed up the testing of new ideas and directions. The gene lists in Tables 3, S1, S2, S3, S4 that compare the values of curvature, topology, geometry, feedback connectivity, and other properties of the networks under study, permit a selection of the best ways to measure lists of genes that impact success of immunotherapy. The conclusion of the analysis presented in this work is that the stability or instability of local network robustness driving changes in feedback connectivity has the largest impact upon prognosis after immunotherapy. The analysis identifies the mutant TP53 gene and its loss of functional protein, resulting in the inability to control cyclin E activity and the resultant abnormalities in copying centrosome numbers accurately as the driving force for this cancer [34, 36].

In conclusion, a network version of the geometric concept of curvature was introduced to model information variability, robustness, and dysregulation of cancer gene networks. Total curvature, thus formulated for HGSOC, was demonstrated to work better in comparison to other standard metrics for the prediction of response to immunotherapy. Network curvature, formulated in this manner as a consistent information passing measure, thus appears to effectively capture global gene signaling dysregulation, and furthermore functions to identify key contributors to signaling dysregulation. Establishing total curvature as a useful clinical biomarker, possibly in combination with FGA (also proposed as a potential biomarker in ovarian cancer [12]), will require larger datasets in order to further quantify and validate these results.

## Code availability

All genetic data and code will be made publicly available upon manuscript publication.

## Acknowledgements

The research of A.T. was funded in part by grants from the Air Force Office of Scientific Research (FA9550-17-1-0435, FA9550-20-1-0029), and NIH grants (R01-AG048769, R21-CA234752). J.D. and A.T. are supported by a grant from the Breast Cancer Research Foundation (BCRF-17-193). D.Z. is supported by the Ovarian Cancer Research Foundation Liz Tilberis Award, and the Department of Defense Ovarian Cancer Research Academy (OC150111). J.R.-F. and B.W. are funded in part by the Breast Cancer Research Foundation and by a grant from the National Institutes of Health/ National Cancer Institute (P50 CA247749 01). The research was supported by the MSK Cancer Center Support Grant/Core Grant (P30 CA008748).

## Author Contributions

R.E. and A.T. developed the mathematical methods, and J.O. developed the bioinformatic analysis. A.L. provided the critical biological and clinical analysis and interpretation. L.N. conceived the project, and contributed to the clinical and biological interpretation of the methodology. J.D. provided insights into interpreting the results and clarifying the technical methods, and D.Z. provided key clinical insights. R.E. wrote the paper, and all authors edited the paper. D.Z., J.R.-F., Y.L., P.S., and B.W. provided the data and assisted in the clinical interpretation of the results. All authors have read and approved the final manuscript.

## Potential Conflict of Interests

D.Z. reports clinical research support to his institution from Astra Zeneca, Plexxikon, and Genentech; and personal/consultancy fees from Merck, Synlogic Therapeutics, GSK, Bristol Myers Squibb, Genentech, Xencor, Memgen, and Agenus. These are all outside of the scope of the submitted work.

J.R.-F. reports receiving personal/consultancy fees from Goldman Sachs, REPARE Therapeutics and Paige.AI, membership of the scientific advisory boards of VolitionRx, REPARE Therapeutics and Paige.AI, membership of the Board of Directors of Grupo Oncoclinicas, and ad hoc membership of the scientific advisory boards of Roche Tissue Diagnostics, Ventana Medical Systems, Novartis, Genentech and InVicro. These are all outside the scope of the submitted work.

B.W. reports ad hoc membership of the advisory board of Repare Therapeutics, outside the scope of the submitted work.

J.D. is a shareholder in PaigeAI. This is outside the scope of the submitted work.

Y.L. reports research funding from AstraZeneca and GSK/Tesaro outside the scope of the submitted work.

None of the other authors report a potential COI.

