## Supplementary Material for "Geometric Network Analysis Provides Prognostic Information in Patients with High Grade Serous Carcinoma of the Ovary Treated with Immune Checkpoint Inhibitors"

### Validation of survival analysis

In this work, curvature is used as a relative measure of network response to immunotherapy in HGS for predicting survival. An independent data set for HGS patients treated with immunotherapy was not available for external validation and the sample size was too small to separate into training and validation sets. We therefore performed three tests for internal validation. $K$-fold cross-validation ($K$-CV) [37] (Supplementary Figure S1) and bootstrap validation (Supplementary Figure S2) were performed to test the validation. In addition, to test if the significant difference in survival found between the high and low curvature groups would likely be observed regardless of the initial gene level data, the CN data was randomly permuted and reassigned amongst the genes. The curvature pipeline was subsequently reperformed with the randomized node weightings and the survival between the two groups defined by low and high curvature according to the 25th percentile of the total curvature value was reassessed in the same manner using the log-rank test. This random process was repeated 500 times with zero out of the 500 trials resulting in a log-rank test statistic greater than the unpermuted sample based test statistic, suggesting that the null hypothesis may fairly be rejected and that total curvature is statistically likely to be picking up on real signals of functional robustness providing a relative measure of overall survival in response to immunotherapy.

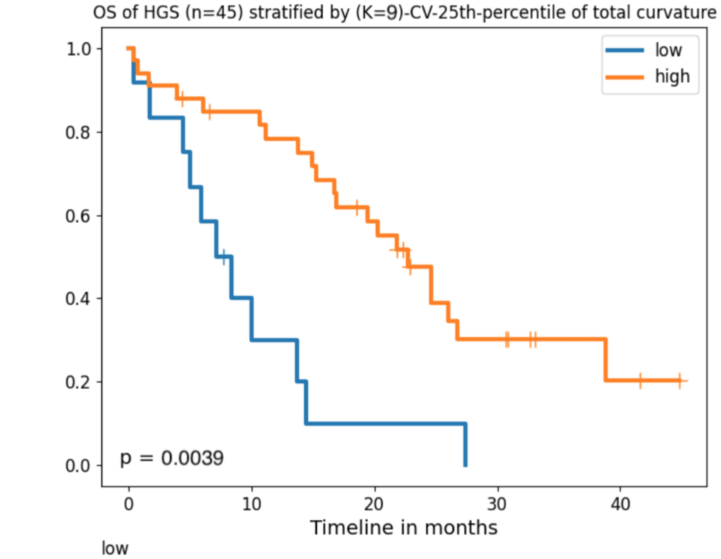


| | | | | | |
|---|---|---|---|---|---|
| **low** | | | | | |
| At risk | 12 | 3 | 1 | 0 | 0 |
| Censored | 0 | 1 | 1 | 1 | 1 |
| Events | 0 | 8 | 10 | 11 | 11 |
| **high** | | | | | |
| At risk | 33 | 26 | 17 | 7 | 2 |
| Censored | 0 | 2 | 3 | 6 | 10 |
| Events | 0 | 5 | 13 | 20 | 21 |

Figure S1: For HGS (n=45), 9-fold cross validated classification Kaplan-Meier curves and p-value determined from the empirical null distribution of the cross-validated log-rank statistic [37]. Briefly, the K-CV was performed as follows: the data set was partitioned into $K$ groups of size $n/K$ where $n$ is the total number of samples (here, $n = 45$, $K = 9$ and $n/K = 5$). The $n/K$ samples in the first partition were removed from the set. The cut for classification was determined to be the 25th percentile of the total curvature from the remaining $n - n/K$ samples. Each of the $n/K$ removed samples was then classified as *low* or *high* curvature by comparing their total curvature to the cutoff. The $n/K$ samples were then returned to the set and the process was repeated for each partition. This process terminated with each sample being classified by $n - n/K$ other samples resulting in two cross-validated high and low curvature groups. KM survival analysis was then performed between the predicted high and low risk groups and a log-rank statistic was computed. The 9-fold CV procedure was repeated 10,000 times. Out of the 10,000 tests, 39 trials resulted in cross-validated log-rank statistic greater than that of original data with test statistic $= 9.499$, resulting in a significance level of 0.0039.

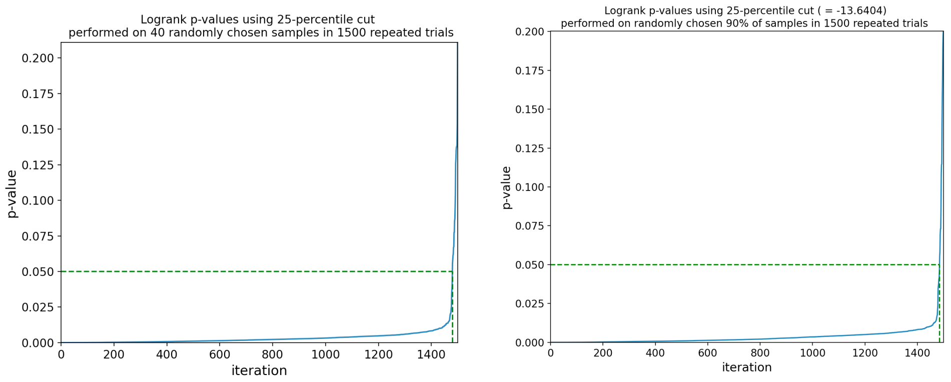


(a) bootstrap cutoff (b) fixed cutoff

Figure S2: HGS (n=45) bootstrapped validation for Kaplan-Meier survival analysis. Bootstrap samples were generated by randomly selecting approximately 90% ($n = 40$) of the HGS cancers. The bootstrap sample was then stratified into high and low curvature groups by either (a) the bootstrap cutoff (the 25th percentile of the bootstrapped set's total curvature values) or (b) the fixed cutoff (the 25th percentile of the original set's total curvature values). In both cases, survival of each bootstrap group was estimated by the Kaplan-Meier analysis and the log-rank p-value was used to test if the survival curves for each group were different. P-values resulting from 1,500 iterations of this process are shown in increasing order by the blue lines. The dotted green lines emphasize the significance level at $\alpha = 0.05$. Out of the 1,500 trials, (a) 20 bootstrap-cutoff trials resulted in $p > 0.05$ and (b) 15 fixed-cutoff trials resulted in $p > 0.05$.

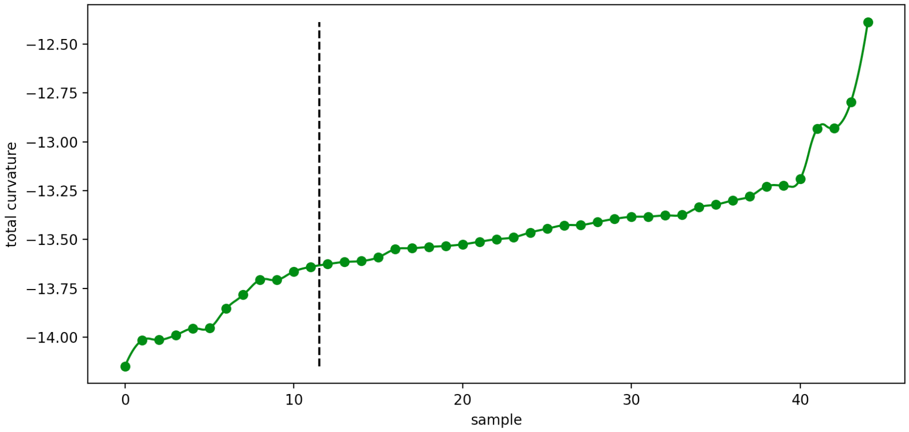


Figure S3: Rationale for selecting 25th-percentile cutpoint. The sorted total curvatures of all samples are shown by green circles. The dotted black line separates the low and high classified samples, where curvature is seen to start increasing slowly by the fitted line.

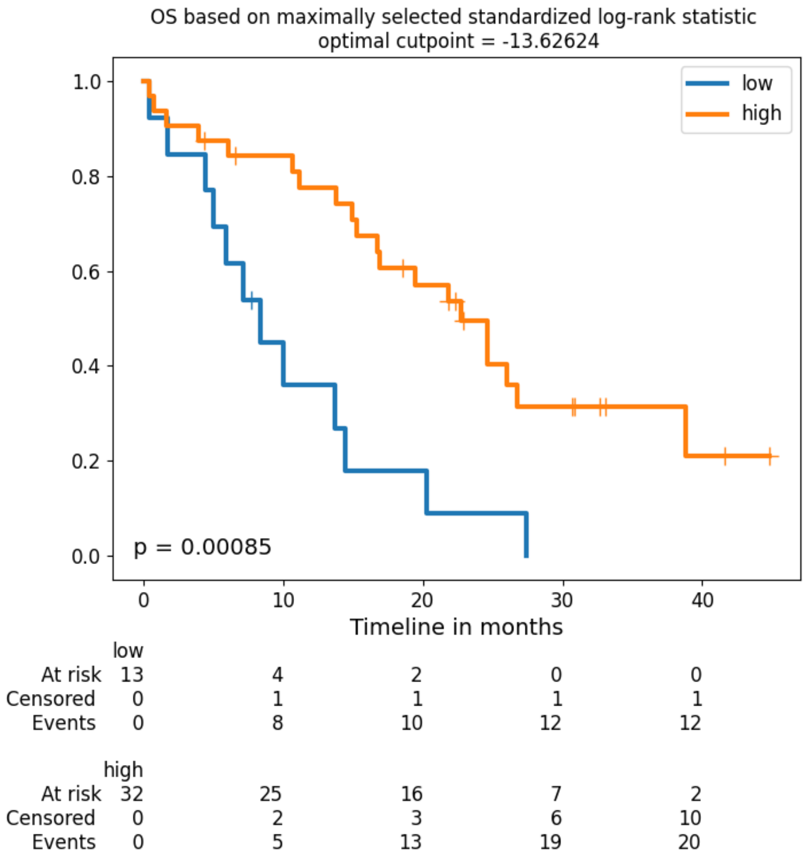


Figure S4: Alternative cutpoint selection. Kaplan-Meier survival analysis using alternative cutpoint for identifying high and low curvature groups with the maximally selected log-rank statistic [31, 32] (cutpoint = -13.62624, requiring 25% minimal proportion) using R's MAXSTAT package. Note that this results in one sample being moved from the high curvature to the low curvature group as found by the 25th percentile of total curvature cutoff.

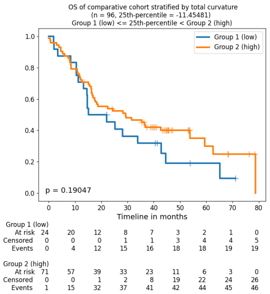


(a) total curvature

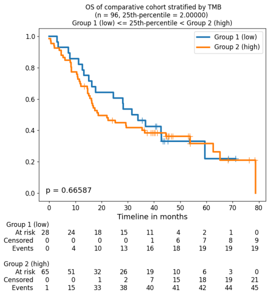


(b) TMB

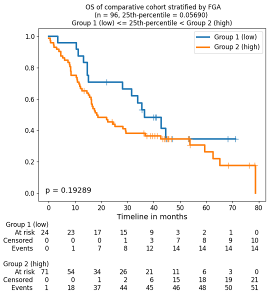


(c) FGA

Figure S5: Survival curves for comparison HGS samples ($n = 96$) stratified low and high groups by the 25th percentile of total curvature and available genomic parameters. Overall survival is defined as the duration from the time of diagnosis to death or time of last follow-up and p-values were derived from the log-rank test.

### Top curvature ranked genes

In this section, we list in Tables S1, S2, S3 the top ranked genes according to the criteria described above in Section 3.2. We list both the greatest positive and negative curvature differences. Finally, in Table S4, we list (in alphabetical order) the key genes found by the curvature analysis using all the comparisons, and then in Table S5, we give the 100 identified candidate genes based on risk listed alphabetically.

| rank | gene | $\Delta\kappa_{OS} > 0$ | gene | $\Delta\kappa_{OS} < 0$ |
|---|---|---|---|---|
| 0 | TP53 | 0.059206 | CTNNB1 | -0.029402 |
| 1 | SMAD3 | 0.055706 | CREBBP | -0.026285 |
| 2 | ATXN1 | 0.028886 | MYC | -0.024026 |
| 3 | EP300 | 0.025355 | PTK2 | -0.023118 |
| 4 | TGFBR1 | 0.017961 | AR | -0.021899 |
| 5 | AKT1 | 0.015810 | SHC1 | -0.018774 |
| 6 | JUN | 0.015342 | SMAD2 | -0.015914 |
| 7 | SRC | 0.013336 | RB1 | -0.015029 |
| 8 | ACTB | 0.013162 | VIM | -0.011736 |
| 9 | PCNA | 0.010664 | PRKCA | -0.010209 |
| 10 | ESR1 | 0.010068 | SMAD4 | -0.009694 |
| 11 | CDKN1A | 0.009641 | MAPK1 | -0.008352 |
| 12 | RAC1 | 0.009197 | GRB2 | -0.008095 |
| 13 | CDKN1B | 0.008439 | SVIL | -0.005490 |
| 14 | PRKCD | 0.008138 | APP | -0.005477 |
| 15 | HSP90AA1 | 0.007070 | SMARCA4 | -0.005419 |
| 16 | CCNE1 | 0.005931 | PARP1 | -0.005321 |
| 17 | STAT1 | 0.005750 | FN1 | -0.005122 |
| 18 | COPS6 | 0.005661 | PIK3R2 | -0.004975 |
| 19 | MAPK14 | 0.005247 | CRMP1 | -0.004917 |
| 20 | MDFI | 0.005216 | MAPK8 | -0.004334 |
| 21 | SMURF1 | 0.005144 | ITGB1 | -0.004226 |
| 22 | CDK5 | 0.004177 | HTT | -0.004059 |
| 23 | ACTN1 | 0.004095 | HSF1 | -0.004010 |
| 24 | YWHAE | 0.003982 | INSR | -0.003442 |
| 25 | DLG4 | 0.003779 | LYN | -0.003200 |
| 26 | C14orf1 | 0.003745 | BTK | -0.003056 |
| 27 | JAK1 | 0.003667 | JAK3 | -0.003018 |
| 28 | PIAS1 | 0.003654 | YAP1 | -0.002875 |
| 29 | FOS | 0.003410 | GSK3B | -0.002822 |
| 30 | PLCG1 | 0.003333 | ATM | -0.002806 |
| 31 | CHD3 | 0.003285 | YWHAQ | -0.002480 |
| 32 | EWSR1 | 0.003279 | BCL2 | -0.002338 |
| 33 | PIK3R1 | 0.003136 | WAS | -0.002229 |
| 34 | NFKBIA | 0.003024 | PPP2R5A | -0.002188 |
| 35 | PML | 0.002953 | MUC1 | -0.002134 |
| 36 | PRNP | 0.002944 | SUV39H1 | -0.002114 |
| 37 | RBPMS | 0.002917 | TGFBR2 | -0.002039 |
| 38 | CCND3 | 0.002851 | ADAM15 | -0.001992 |
| 39 | RUNX2 | 0.002838 | MAGEA11 | -0.001903 |
| 40 | MAP2K1 | 0.002709 | POU2F1 | -0.001759 |
| 41 | PSEN1 | 0.002696 | SYN1 | -0.001755 |
| 42 | BRCA1 | 0.002660 | PRKCG | -0.001745 |
| 43 | XPO1 | 0.002648 | RGS2 | -0.001736 |
| 44 | DVL2 | 0.002554 | DNM2 | -0.001634 |
| 45 | CRK | 0.002550 | PAK1 | -0.001629 |
| 46 | TRAF6 | 0.002531 | FEZ1 | -0.001536 |
| 47 | MCM7 | 0.002298 | JAK2 | -0.001488 |
| 48 | NEDD4 | 0.002287 | UPF1 | -0.001487 |
| 49 | RAD51 | 0.002286 | MDM2 | -0.001449 |

Table S1: Comparing average scalar curvature based on overall survival (OS). Top 50 genes ranked by positive ($\Delta\kappa_{OS} > 0$) and negative ($\Delta\kappa_{OS} < 0$) difference in average scalar curvature between alive ($n = 13$) and dead ($n = 32$) cohorts [illegible]

| rank | gene | $\Delta\kappa_{PM} > 0$ | gene | $\Delta\kappa_{PM} < 0$ |
|---:|---|---:|---|---:|
| 0 | TP53 | 0.077029 | ESR1 | -0.036440 |
| 1 | GRB2 | 0.052007 | SMAD3 | -0.021618 |
| 2 | ATXN1 | 0.047675 | CDKN1A | -0.020347 |
| 3 | PRKCA | 0.042002 | JUN | -0.015349 |
| 4 | CREBBP | 0.026753 | SRC | -0.013494 |
| 5 | SMAD2 | 0.019927 | LYN | -0.011455 |
| 6 | AKT1 | 0.017599 | MDFI | -0.010594 |
| 7 | TGFBR1 | 0.012515 | EGFR | -0.008569 |
| 8 | SMAD4 | 0.011964 | MAPK14 | -0.008072 |
| 9 | CTNNB1 | 0.010307 | CCNE1 | -0.007612 |
| 10 | EWSR1 | 0.010187 | GSK3B | -0.006226 |
| 11 | VIM | 0.009960 | RB1 | -0.005719 |
| 12 | EP300 | 0.009048 | RUNX2 | -0.005594 |
| 13 | APP | 0.007988 | CCND3 | -0.005412 |
| 14 | HSP90AA1 | 0.007883 | MYOC | -0.005080 |
| 15 | MYC | 0.007121 | JAK2 | -0.004539 |
| 16 | PTK2 | 0.005757 | CDKN1B | -0.004367 |
| 17 | HGS | 0.005709 | DAXX | -0.004313 |
| 18 | YWHAE | 0.005597 | RBPMS | -0.004050 |
| 19 | DLG4 | 0.005540 | PIK3R2 | -0.004026 |
| 20 | SVIL | 0.005213 | MAP3K7 | -0.003988 |
| 21 | COIL | 0.005048 | PRKCD | -0.003983 |
| 22 | PIK3R1 | 0.004457 | MDM2 | -0.003608 |
| 23 | ITGB1 | 0.004411 | NCOA2 | -0.003404 |
| 24 | STAT1 | 0.004319 | RAF1 | -0.003399 |
| 25 | SLC9A3R1 | 0.004226 | RAC1 | -0.003303 |
| 26 | MAPK1 | 0.004204 | AKT2 | -0.003245 |
| 27 | ABL1 | 0.003941 | PLCG1 | -0.002819 |
| 28 | CDK5 | 0.003806 | HRAS | -0.002807 |
| 29 | BRCA1 | 0.003210 | IGF1R | -0.002795 |
| 30 | ACTN1 | 0.003170 | EZR | -0.002769 |
| 31 | RANBP9 | 0.003063 | MLLT4 | -0.002732 |
| 32 | PRKCG | 0.002898 | PRKCB | -0.002682 |
| 33 | GFI1B | 0.002779 | POU2F1 | -0.002679 |
| 34 | TLE1 | 0.002757 | FASLG | -0.002679 |
| 35 | CHD3 | 0.002688 | JAK1 | -0.002676 |
| 36 | CCND1 | 0.002670 | GNAI1 | -0.002552 |
| 37 | SYK | 0.002652 | PCNA | -0.002510 |
| 38 | MAPK8 | 0.002550 | PLG | -0.002244 |
| 39 | DVL2 | 0.002504 | FN1 | -0.002154 |
| 40 | YAP1 | 0.002367 | XPO1 | -0.002072 |
| 41 | BCL2 | 0.002288 | ACTB | -0.002056 |
| 42 | HTT | 0.002270 | CDC42 | -0.002046 |
| 43 | POLR2A | 0.002198 | SHC1 | -0.001900 |
| 44 | TRIP13 | 0.002189 | SUMO4 | -0.001896 |
| 45 | ATM | 0.002159 | JAK3 | -0.001888 |
| 46 | CRMP1 | 0.002127 | COPS5 | -0.001875 |
| 47 | ACTG1 | 0.002125 | CD247 | -0.001809 |
| 48 | C14orf1 | 0.002115 | PIK3CA | -0.001768 |
| 49 | SUMO2 | 0.001974 | RXRG | -0.001731 |

Table S2: Changes in average scalar curvature based on sample type (primary (P) vs metastasis (M)). Top 50 genes ranked by positive ($\Delta\kappa_{PM} > 0$) and negative ($\Delta\kappa_{PM} < 0$) difference in average scalar curvature between P ($n = 13$) [illegible]

| rank | gene | $\Delta\kappa_{ref} > 0$ | gene | $\Delta\kappa_{ref} < 0$ |
|---:|---|---:|---|---:|
| 0 | TP53 | 0.143999 | SRC | -0.070044 |
| 1 | EP300 | 0.089752 | ATXN1 | -0.041866 |
| 2 | AR | 0.055047 | PTK2 | -0.040546 |
| 3 | TGFBR1 | 0.050407 | MYC | -0.032654 |
| 4 | MAPK1 | 0.041837 | LYN | -0.026977 |
| 5 | ESR1 | 0.025883 | SHC1 | -0.021819 |
| 6 | PIK3R1 | 0.024060 | GSK3B | -0.017265 |
| 7 | SMAD3 | 0.023103 | JUN | -0.015936 |
| 8 | RB1 | 0.021967 | PCNA | -0.015054 |
| 9 | EWSR1 | 0.020344 | PLCG1 | -0.014421 |
| 10 | SMAD4 | 0.019587 | MAPK14 | -0.014055 |
| 11 | SMAD2 | 0.019196 | CDKN1A | -0.011164 |
| 12 | CREBBP | 0.018241 | YWHAQ | -0.008725 |
| 13 | ABL1 | 0.016583 | MDM2 | -0.007339 |
| 14 | CSNK2A2 | 0.012132 | NCOA2 | -0.007238 |
| 15 | DLG4 | 0.011312 | CCNE1 | -0.006786 |
| 16 | CTNNB1 | 0.010916 | CDKN1B | -0.006716 |
| 17 | BRCA1 | 0.010490 | HCK | -0.006688 |
| 18 | EGFR | 0.009949 | PAK1 | -0.006402 |
| 19 | YWHAE | 0.009881 | DAXX | -0.006289 |
| 20 | GFI1B | 0.009811 | HSF1 | -0.006140 |
| 21 | ACTB | 0.009696 | BCL2L1 | -0.005700 |
| 22 | RAC1 | 0.009329 | RBL1 | -0.005566 |
| 23 | MAGEA11 | 0.009010 | MDFI | -0.005182 |
| 24 | BTK | 0.008727 | STAT1 | -0.004959 |
| 25 | XRCC6 | 0.007943 | ACVR1 | -0.004607 |
| 26 | UBB | 0.007720 | SUMO1 | -0.004323 |
| 27 | AKT1 | 0.007642 | RANBP9 | -0.004323 |
| 28 | CHD3 | 0.007446 | COPS5 | -0.004273 |
| 29 | TLE1 | 0.007259 | CDK5 | -0.004024 |
| 30 | SAT1 | 0.006644 | MYOC | -0.003970 |
| 31 | JAK2 | 0.006454 | FN1 | -0.003823 |
| 32 | DVL2 | 0.006230 | PARP1 | -0.003753 |
| 33 | SYK | 0.006148 | CDK4 | -0.003741 |
| 34 | NOTCH1 | 0.005997 | CCND3 | -0.003324 |
| 35 | POLR2A | 0.005812 | COPS6 | -0.003277 |
| 36 | NCOR1 | 0.005568 | SMURF1 | -0.003160 |
| 37 | HSP90AA1 | 0.005143 | HIPK2 | -0.003070 |
| 38 | INSR | 0.005123 | POU2F1 | -0.003066 |
| 39 | CRK | 0.004940 | XPO1 | -0.003044 |
| 40 | PRKCB | 0.004893 | TGM2 | -0.003020 |
| 41 | BCAR1 | 0.004520 | FGFR1 | -0.002781 |
| 42 | HTT | 0.004499 | PRNP | -0.002770 |
| 43 | BCL2 | 0.004241 | MUC1 | -0.002713 |
| 44 | SH3KBP1 | 0.003904 | TRAF6 | -0.002684 |
| 45 | UTP14A | 0.003815 | YAP1 | -0.002674 |
| 46 | PRKCD | 0.003753 | MCM2 | -0.002534 |
| 47 | RASA1 | 0.003679 | RUNX2 | -0.002534 |
| 48 | RARA | 0.003616 | PRSS23 | -0.002533 |
| 49 | FXR2 | 0.003610 | NINL | -0.002514 |

Table S3: Top 50 genes ranked by positive ($\Delta\kappa_{ref} > 0$) and negative ($\Delta\kappa_{ref} < 0$) difference between average HGS scalar curvature ($n = 45$) $\kappa_{HGS}$ and the scalar curvature of the reference topology $\kappa_{top}$. $\Delta\kappa_{ref} = \kappa_{HGS} - \kappa_{top}$.

| | | | | |
|---|---|---|---|---|
| ABL1 | CTNNB1 | MAP2K1 | PRKCE | SYK |
| ACTB | DAXX | MAP3K7 | PRKCG | SYN1 |
| ACTG1 | DLG4 | MAPK1 | PRNP | TGFBR1 |
| ACTN1 | DNM2 | MAPK14 | PRSS23 | TGFBR2 |
| ACVR1 | DVL2 | MAPK8 | PSEN1 | TGM2 |
| ADAM15 | EGFR | MCM2 | PTK2 | TLE1 |
| AKT1 | EIF2AK2 | MCM7 | RAC1 | TP53 |
| AKT2 | EP300 | MDFI | RAD51 | TRAF6 |
| APP | ESR1 | MDM2 | RAF1 | TRIP13 |
| AR | EWSR1 | MLLT4 | RANBP9 | UBB |
| ARRB2 | EZR | MUC1 | RARA | UPF1 |
| ATM | FASLG | MYC | RASA1 | UTP14A |
| ATXN1 | FEZ1 | MYOC | RB1 | VIM |
| AXIN1 | FGFR1 | NCOA2 | RBL1 | WAS |
| BCAR1 | FN1 | NCOR1 | RBPMS | XPO1 |
| BCL2 | FOS | NEDD4 | RGS2 | XRCC6 |
| BCL2L1 | FXR2 | NFKBIA | RHOA | YAP1 |
| BRCA1 | GFI1B | NINL | RPA1 | YWHAE |
| BTK | GNAI1 | NOTCH1 | RUNX2 | YWHAQ |
| C14orf1 | GRB2 | NR3C1 | RXRG | |
| CASP8 | GSK3B | NTRK1 | SAT1 | |
| CCND1 | HCK | PAK1 | SH3KBP1 | |
| CCND3 | HDAC3 | PARP1 | SHC1 | |
| CCNE1 | HGS | PCNA | SLC9A3R1 | |
| CD247 | HIPK2 | PDPK1 | SMAD2 | |
| CDC42 | HRAS | PIAS1 | SMAD3 | |
| CDK4 | HSF1 | PIK3CA | SMAD4 | |
| CDK5 | HSP90AA1 | PIK3R1 | SMAD7 | |
| CDKN1A | HTT | PIK3R2 | SMARCA4 | |
| CDKN1B | IGF1R | PLCG1 | SMURF1 | |
| CHD3 | INSR | PLG | SNAPIN | |
| COIL | ITGB1 | PML | SRC | |
| COPS5 | JAK1 | POLR2A | STAT1 | |
| COPS6 | JAK2 | POU2F1 | SUMO1 | |
| CREBBP | JAK3 | PPP2R5A | SUMO2 | |
| CRK | JUN | PRKCA | SUMO4 | |
| CRMP1 | LYN | PRKCB | SUV39H1 | |
| CSNK2A2 | MAGEA11 | PRKCD | SVIL | |

Table S4: Intersection of 171 top ranked candidate genes listed alphabetically.

| | | | | |
|---|---|---|---|---|
| ACTB | ACVR1 | ADAM15 | AKT1 | APP |
| AR | ARRB2 | ATXN1 | AXIN1 | BCL2 |
| BRCA1 | BTK | CASP8 | CD247 | CDC42 |
| CDK5 | CDKN1A | CHD3 | COIL | COPS6 |
| CREBBP | CRK | CRMP1 | CSNK2A2 | CTNNB1 |
| DLG4 | DVL2 | EGFR | EIF2AK2 | EP300 |
| ESR1 | EWSR1 | FASLG | FGFR1 | FN1 |
| FXR2 | GNAI1 | GRB2 | GSK3B | HDAC3 |
| HGS | HIPK2 | HRAS | HSF1 | HSP90AA1 |
| HTT | JAK1 | JUN | LYN | MAGEA11 |
| MAPK1 | MAPK14 | MDM2 | MUC1 | MYC |
| MYOC | NCOR1 | NR3C1 | NTRK1 | PAK1 |
| PARP1 | PCNA | PDPK1 | PIK3R1 | PIK3R2 |
| PLCG1 | POLR2A | POU2F1 | PPP2R5A | PRKCA |
| PRKCD | PRKCE | PTK2 | RAC1 | RAF1 |
| RANBP9 | RASA1 | RB1 | RHOA | RPA1 |
| SHC1 | SMAD2 | SMAD3 | SMAD4 | SMAD7 |
| SMARCA4 | SMURF1 | SNAPIN | SRC | STAT1 |
| SUMO1 | SUMO4 | TGFBR1 | TP53 | UBB |
| VIM | XPO1 | XRCC6 | YWHAE | YWHAQ |

Table S5: 100 identified candidate genes based on risk listed alphabetically

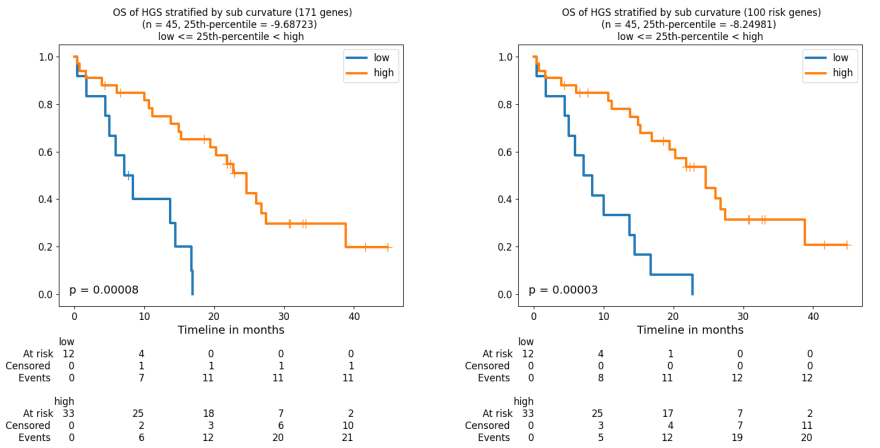


(a) (b)

Figure S6: Survival curves for HGS patients (n=45) based on the 25th percentile of sub-curvature over the following subsets of genes: (a) all 171 curvature identified genes (Supplementary Table S4) (b) 100 risk identified genes (Supplementary Table S5)

### Sub-curvature survival analysis

The Kaplan-Meier analysis suggests that total curvature as a network measure of functional robustness is a more effective indicator of survival in HGSOC treated with ICIs than other genomic parameters. However, we expected that not all genes in the network were necessary to predict survival. We therefore defined the *sub-curvature* $\kappa_g$ for any subset of genes $g$ to be the sum of scalar (node) curvatures over all genes in the given subset:

$$\kappa_g = \sum j \in g\kappa_j. \tag{16}$$

We then repeated the survival analysis replacing total curvature with sub-curvature using the following two subsets of genes: (1) the *curvature identified genes* consisting of the 171 unique genes top-ranked by the various network curvature criteria (Supplementary Tables S1,S2,S3,3) listed alphabetically in Supplementary Table S4 and (2) the *risk identified genes* consisting of the top-ranked 100 genes by the curvature risk criterion (Table 3) listed alphabetically in Supplementary Table S5. Survival curves based on the 25th-percentile of sub-curvature values for each of these subsets are shown in Figure S6. In both cases, the p-value (171 curvature identified genes: $p = 0.00008$; 100 risk identified genes: $p = 0.00003$) is about 1 order of magnitude smaller as compared to the total curvature, suggesting that the curvature methodology identifies key genes pertaining to survival.

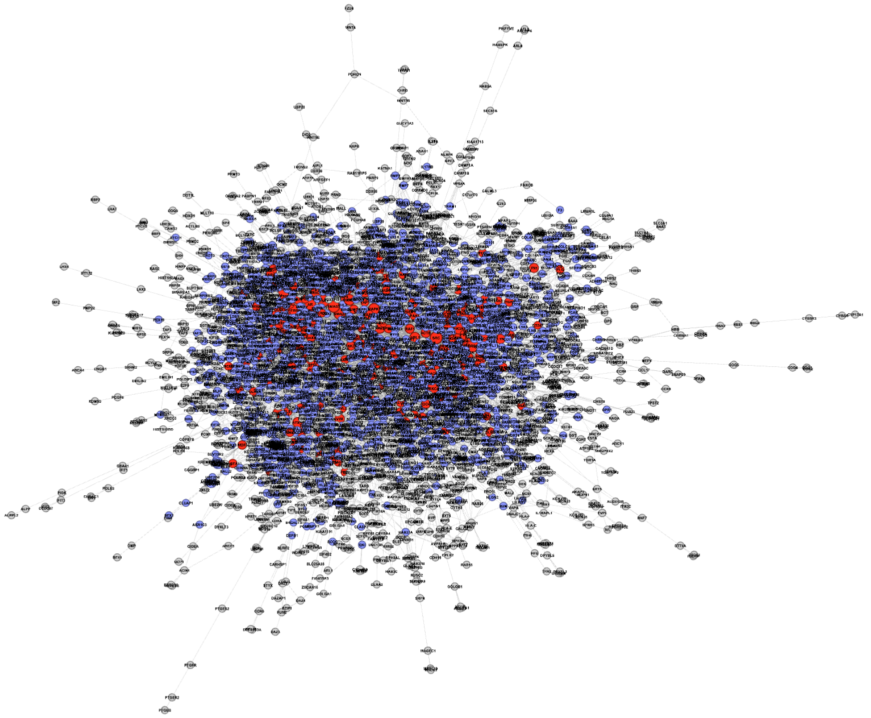

Figure S7: Full resulting network taken as the largest connected component of the intersection between the HPRD and data (3,489 nodes, 9,710 edges, average degree = 5.57) is shown in ForceAtlas 2 layout using Gephi [38]. Top 171 identified genes are shown in red. Neighbors of the identified genes are shown in blue. Node size is scaled by degree.

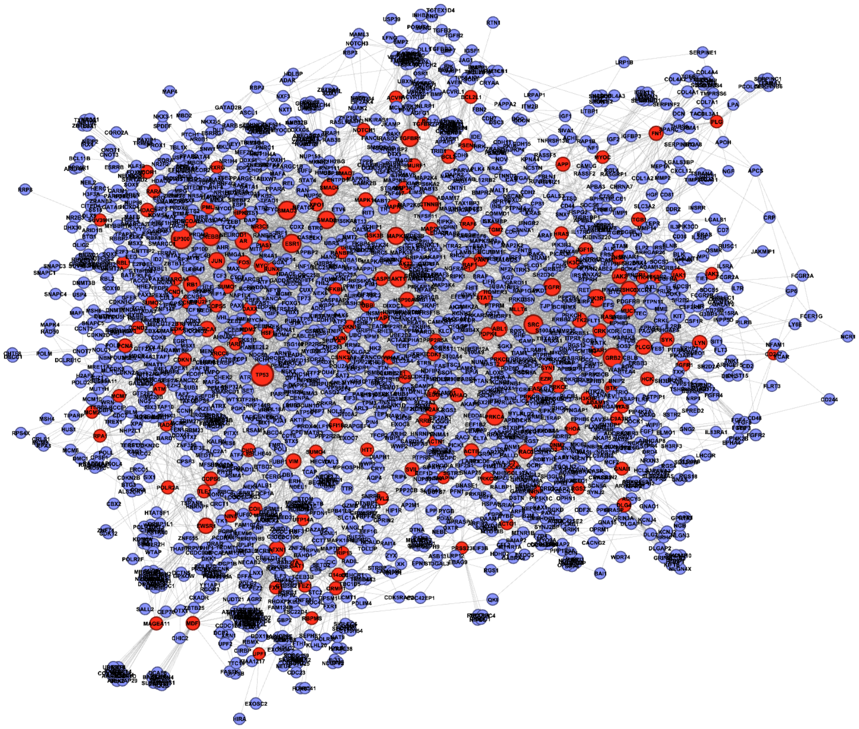

Figure S8: Subnetwork consisting of top 171 identified genes (red) and their neighbors (blue) has 2,084 nodes in total, 7,359 edges and average degree = 7.06. Nodes are scaled by degree and the configuration was generated using Gephi's ForceAtlas 2 layout [38].

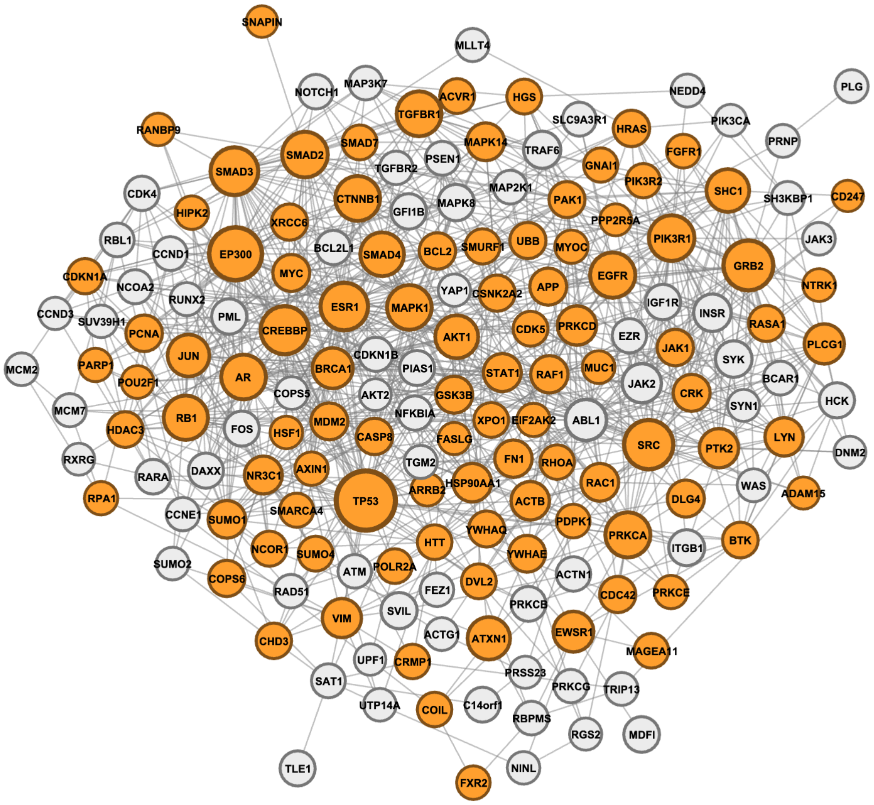


Figure S9: Subnetwork consisting of top 171 identified genes. The 100 risk associated genes are shown in orange. Nodes are scaled by degree and the configuration was generated using Gephi's ForceAtlas 2 layout [38].
